# Parallel duplication and loss of aquaporin-coding genes during the ‘out of the sea’ transition as potential key drivers of animal terrestrialization

**DOI:** 10.1101/2022.07.25.501387

**Authors:** Gemma I. Martínez-Redondo, Carolina Simón Guerrero, Leandro Aristide, Pau Balart-García, Vanina Tonzo, Rosa Fernández

## Abstract

One of the most important physiological challenges animals had to overcome during terrestrialization (i.e., the transition from sea to land) is water loss, which alters their osmotic and hydric homeostasis. Aquaporins are a superfamily of membrane water transporters heavily involved in osmoregulatory processes. Their diversity and evolutionary dynamics in most animal lineages remain unknown, hampering our understanding of their role in marine-terrestrial transitions. Here, we interrogated aquaporin gene repertoire evolution across the main terrestrial animal lineages. We annotated aquaporin-coding genes in genomic data from 458 species from 7 animal phyla where terrestrialization episodes occurred. We then explored aquaporin gene evolutionary dynamics to assess differences between terrestrial and aquatic species through phylogenomics and phylogenetic comparative methods. Our results revealed parallel aquaporin-coding gene duplications in aquaporins during the transition from marine to non-marine environments (e.g., brackish, freshwater and terrestrial), rather than from aquatic to terrestrial ones, with some notable duplications in ancient lineages. Contrarily, we also recovered a significantly lower number of superaquaporin genes in terrestrial arthropods, suggesting that more efficient oxygen homeostasis in land arthropods might be linked to a reduction in this type of aquaporins. Our results thus indicate that aquaporin-coding gene duplication and loss might have been one of the key steps towards the evolution of osmoregulation across animals, facilitating the ‘out of the sea’ transition and ultimately the colonisation of land.

## Introduction

The ecological transition from water to land (i.e. terrestrialization) is one of the most remarkable episodes that shaped the evolution of biodiversity as we know it today. Throughout the eukaryote tree of life, living beings have shifted their habitat from water to land multiple times (Jamy et al., 2022). Among them, animals have successfully terrestrialized independently in 10 different phyla: chordates, nematodes, nematomorphs, tardigrades, arthropods, onychophorans, platyhelminthes, molluscs, annelids, and nemerteans. Within most phyla, more than one independent habitat shift has occurred, ranging from only one terrestrialization event in the onychophorans to more than 30 in nematodes (Holterman, Schratzberger, & Helder, 2019), at different times along Earth’s history. The most studied terrestrialization event is, beyond doubt, the one that gave rise to terrestrial tetrapods during the Devonian period from euryhaline aquatic ancestors (Goedert et al., 2018; Qvarnström, Szrek, Ahlberg, & Niedźwiedzki, 2018). Nevertheless, the most ancient terrestrialization events occurred in arthropods (arachnids, myriapods and hexapods) from marine ancestors during the late Cambrian-early Ordovician (Rota-Stabelli, Daley, & Pisani, 2013). More than 4 additional independent terrestrialization events have occurred in the following millions of years within Pancrustacea (isopods, amphipods and anomura and brachyura decapods) (Sharma, 2017; Watson-Zink, 2021). Other terrestrialization events that have recently gained interest are the ones in molluscs and annelids. In the case of molluscs, gastropods have independently colonised land more than 9 times (Kameda & Kato, 2011), with the latest estimates suggesting up to 30 times (Vermeij & Watson-Zink, 2022), either following a freshwater route or through marginal environments such as brackish water (Krug et al., 2022; van Straalen, 2021; Vermeij & Watson-Zink, 2022). Among them, the most successful terrestrial gastropod lineage is the Stylommatophora, an order of air-breathing pulmonate land gastropods that contains most living snails and slugs. Regarding annelids, earthworms (Crassiclitellata) derive from a single terrestrialization event, while other events have occurred within Hirudinida and Enchytraeidae (Erséus et al., 2020; van Straalen, 2021). Another example is that of onychophorans, or velvet worms, which are the only entirely terrestrial animal phylum. This lineage closely related to arthropods has been hypothesised to have a marine origin based on the similar osmotic pressure of their haemolymph to terrestrial arthropods (Little, 1990). All these independent colonisation events (Fig 1A) make animal terrestrialization such a complex and intriguing area of study, especially since the genomic underpinnings facilitating these transitions remain highly unexplored.

**Figure 1.** Terrestrialization and aquaporins. **A**. Schematic representation of the main terrestrialization events in the phyla included in this study and their habitat of origin. Arthropods and onychophorans have mostly colonised land from a marine ancestor; some terrestrial gastropods and Crassiclitellata annelids (earthworms) from a freshwater ancestor; while tetrapods and other terrestrial molluscs (such as Stylommatophora) have a euryhaline and mangrove/intertidal origin, respectively. In the case of hexapods, their terrestrialization has been suggested to have been mediated through an intermediate subterranean state, represented by a cave in the figure, due to their relatedness with the subterranean aquatic Remipedia. **B**. Aquaporin tertiary and quaternary structure and representation of transmembrane domains adapted from(Nesverova & Törnroth-Horsefield, 2019). Water molecules are represented in red, and the two half-helices formed by loops B and E are in yellow. **C**. Schematic representation of pore size and structure changes allow different molecules to pass through. Molecules are ordered from the smallest to the largest (water, ammonia, and glycerol, respectively) from left to right. **D**. General aquaporin phylogeny based on Abascal et al., 2014.

Terrestrialization required many complex physiological changes in different biological processes, such as respiration, locomotion or reproduction, that allowed them to overcome the environmental barriers and adapt to life on land. One of the most important challenges species faced during this transition was avoiding water loss, which alters their osmotic and hydric homeostasis. Most marine animals are osmoconformers, which means that they cannot regulate their internal osmotic pressure under environmental shifts. When marine animals transitioned to euryhaline or freshwater habitats, they faced osmotic stress due to the differences in the ionic composition between their extracellular fluids and the environment. The development of the ability to osmoregulate, i.e. to maintain the osmotic pressure of their extracellular fluids (Rankin & Davenport, 1981), allowed them to overcome this osmotic shock. Accordingly, terrestrial animals have acquired several physiological adaptive mechanisms to regulate their internal osmotic balance and water loss. Such is the case of closing spiracles in arthropods with tracheae or the pneumostome in terrestrial pulmonate gastropods to regulate gas exchange, the evolution of a kidney structure and non-ammonia excretion in tetrapods, or the development of Malpighian tubules in terrestrial arthropods for regulating osmotic pressure.

At the molecular level, animals can modify the osmotic composition of their fluids by regulating the transport of ions or water through biological membranes (Takvam, Wood, Kryvi, & Nilsen, 2021; Uchiyama & Konno, 2006). Previous experiments have shown that cells of some of these osmoregulatory organs express protein channels called aquaporins in their membranes (Cabrero et al., 2020; Finn, Chauvigné, Hlidberg, Cutler, & Cerdà, 2014;Kaufmann et al., 2005; Spring, Robichaux, & Hamlin, 2009; Wittekindt & Dietl, 2019; Yi et al.,2011). Aquaporins are a superfamily of small membrane intrinsic proteins (24-36kDa) that are heavily involved in osmoregulatory processes by forming pores in the cell membrane that facilitate the transport of water and other small molecules (Abascal, Irisarri, & Zardoya, 2014;Kruse, Uehlein, & Kaldenhoff, 2006). Structurally, the basic units of aquaporins are monomers characterised by six transmembrane domains that surround a narrow pore, and two selectivity filters: two NPA (Asn-Pro-Ala) motifs, responsible for proton exclusion, and an aromatic/arginine (ar/R) region that determines the pore size (Fig 1B). Variations in these two selectivity filters slightly change the pores’ size, structure and hydrophobicity, and ultimately determine different subtypes of aquaporins and their ability to transport molecules different from water (Fig 1C) (de Maré, Venskutonytė, Eltschkner, de Groot, & Lindkvist-Petersson,2020). In the membrane, four aquaporin monomers form a tetramer, creating a fifth pore (Kreida & Törnroth-Horsefield, 2015). This general structure is conserved between bacteria and eukaryotic aquaporins, as well as between different subtypes of aquaporins, despite the low sequence similarity of these proteins (less than 30%) (Kruse et al., 2006).

First phylogenetic analysis of aquaporins across all organisms, together with the permeability differences of aquaporins, allowed their classification into two main subtypes: aquaporins that transport water (Aqps), and aquaglyceroporins that are also permeable to glycerol and other molecules (Glps). Further studies showed that both are also permeable to other molecules (CO_2_, H_2_O_2_ or ammonia, among others) (Finn & Cerdà, 2015). In animals, Aqps can be further divided into three subtypes: classical aquaporins, aquaammoniaporins, and superaquaporins (the name that we will use hereafter instead of unorthodox aquaporins as suggested by (Benga, 2012)), whose origin dates back to the base of the eukaryotic tree (Fig 1D) (Abascal et al., 2014).

Previous studies have analysed the repertoire of aquaporin-coding genes (Aqp-coding genes hereafter) in selected animal species (e.g., *Danio rerio* (Tingaud-Sequeira et al., 2010), *Caenorhabditis elegans (Huang, Lamitina, Agre, & Strange, 2007), Milnesium tardigradum (Grohme et al., 2013)*, blood-feeding arthropods (Benoit
 et al., 2014), dust mites (Peng et al., 2018) or the salmon louse and its host (Stavang et al., 2015), to name a few). Meanwhile, others have focused on the diversity and evolution of animal aquaporins in groups of human interest, such as vertebrate (Finn et al., 2014) and hemipteran aquaporins (Van Ekert et al., 2016), or insect entomoglyceroporins (Finn, Chauvigné, Stavang, Belles, & Cerdà, 2015). Recent studies have leveraged the newly available transcriptomic and genomic resources to expand taxon sampling and dig deeper into aquaporin (or aquaglyceroporin) diversity in groups of animals that are not widely studied, such as copepods and crustaceans (Catalán-García et al., 2021), gastropods (Colgan & Santos,2018), annelids (Mucciolo et al., 2021), and non-vertebrate deuterostomes (Yilmaz et al.,2020). However, the whole aquaporin gene repertoire in other animal groups, such as molluscs and less studied orders of arthropods (i.e. chelicerates other than spiders, scorpions and mites), has not yet been explored, let alone in a comparative framework within the context of animal terrestrialization. Only a couple of previous studies have explored the potential key role of aquaporin gene duplications in the adaptation of vertebrates to life on land, with the identification of 3 tetrapod-exclusive classical aquaporins functionally related to water absorption mechanisms (Finn et al., 2014), and some signals of positive selection in semiterrestrial mudskippers’ aquaporins (Lorente-Martínez, Agorreta, Torres-Sánchez, & Mauro, 2018). Nevertheless, even though the importance of gene duplication in the origin of new functions and driving adaptation has long been reported (Kondrashov, 2012), the potential role of aquaporin gene duplication and loss in the terrestrialization of lineages other than tetrapods has not yet been explored.

Here, we test the hypothesis that gene duplication and loss in Aqp-coding genes may have played a key role in animal terrestrialization and facilitated marine-to-land transition in other phyla beyond chordates. To show this, we mined and annotated more than 6,300 Aqp-coding gene sequences from 458 genomic and transcriptomic data sets, maximising taxon sampling to ensure a good representation of most animal lineages with terrestrial representatives. To assess whether aquaporins may have had a role during animal terrestrialization, we explored aquaporin gene evolutionary dynamics to assess differences between terrestrial and aquatic species through phylogenomics and phylogenetic comparative methods. We interpreted and discussed our results in the context of animal terrestrialization.

## Materials and methods

### Obtaining genomic and transcriptomic data and habitat assignment

We analysed the aquaporin gene repertoire of 485 species from 7 animal phyla: Arthropoda, Onychophora, Tardigrada, Nematoda, Mollusca, Annelida and Vertebrata. The full list of species included and the source they were obtained from is available at Supp. Data 1.

Data from the remaining 415 invertebrate species were obtained from MATEdb (Fernández et al., 2022), which provides standard comparable proteomes for phylogenomic studies. We downloaded the gene annotation of those species (longest isoform for each gene), and their taxonomic and habitat information. Vertebrate, *M. tardigradum*, and *C. elegans* aquaporins were directly obtained from the literature (Finn et al., 2014; Grohme et al., 2013; Huang et al., 2007) respectively). The only exceptions were the aquaporins from the amphibian *Leptobrachium leishanense*, for which we downloaded the genome annotation (Cunningham et al., 2021). In the case of annelids, (Mucciolo et al., 2021) provided a fairly good representation of aquaporins across Pleistoannelida, the most abundant clade within annelids. However, as noted by the authors, the methodology followed could have underestimated the identification of aquaporins due to the incompleteness or low quality of the source databases’ data. For instance, An example of this is *Eisenia andrei*, for which only one Aqp was identified. To verify this information, we downloaded the genome annotation of *E. andrei* (Shao et al., 2020) to annotate its Aqp-coding genes ourselves as described below. We identified a total of 27 Aqp-coding genes, in contrast with the one reported by (Mucciolo et al., 2021).

Habitat information for species was obtained from the World Register of Marine Species (WoRMS) database (WoRMS Editorial Board 2022). A species was considered to be aquatic when it spent at least one stage of its life cycle in an aquatic environment.

### Annotating genes encoding for aquaporins in downloaded proteomes

From the whole set of proteomes, genes encoding for aquaporins were annotated using BITACORA version 1.3 (Vizueta, Sánchez-Gracia, & Rozas, 2020) in “protein mode”. Briefly, BITACORA combines BLAST and HMMER (Eddy, 2011) searches against a custom curated database of genes belonging to the gene family of interest. Due to the conservation of aquaporins across all animals, our aquaporin curated database contained a list of 123 metazoan aquaporin sequences that have been experimentally confirmed in previous studies (Supp. Data 2). To benchmark this methodology, we used *Drosophila melanogaster* to compare the number of aquaporins identified using BITACORA with the number of aquaporins annotated in its genome, obtaining the same result.

### Aquaporin-coding genes nomenclature

Aquaporin-coding genes nomenclature between animal phyla is challenging due to the lack of consistency in the different studies (Benoit et al., 2014; Kosicka, Grobys, Kmita,Lesicki, & Pieńkowska, 2016; Kosicka, Lesicki, & Pieńkowska, 2020; Mucciolo et al., 2021;Stavang et al., 2015; Yilmaz et al., 2020). This conflict worsens when considering papers on the identification of aquaporins of isolated species without taking their evolutionary history into account (Grohme et al., 2013; Huang et al., 2007). Here, we follow the system used by (Mucciolo et al., 2021) and (Kosicka et al., 2020) but taking into account the gene nomenclature rules, and name each of the aquaporin subtypes as “AqpX-like”, being X the lowest nonzero number of the vertebrate aquaporin coding-gene that forms part of it. Thus, we define the four subtypes as Aqp3-like (aquaglyceroporins), Aqp1-like (classical aquaporins), Aqp8-like (aquaammoniaporins) and Aqp11-like (superaquaporins), and follow this nomenclature hereafter.

### Phylogenetic tree inference of aquaporin-coding genes

To identify which aquaporins belonged to which subtype (i.e., Aqp1-like, Aqp3-like, Aqp8-like or Aqp11-like) for posterior analyses, we first independently classified aquaporins of all animal species included in this study, except for the ones that had already been classified in previous publications (Finn et al., 2014; Mucciolo et al., 2021). We based this classification on the description of aquaporin phylogenetic clusters and their relative position in the phylogenetic tree to the already classified sequences, which were considered as a backbone.

For arthropods, onychophorans and molluscs (i.e. lineages whose aquaporins we identified de novo in this study) we manually curated the classification by checking unclustered aquaporin sequences and long branches that could represent outliers using BLASTP against the non-redundant NCBI database. This was done to reclassify identified aquaporins that had wrongly fallen into the wrong subtype and remove contaminants from species outside of the studied phylum that had escaped filtering in the original data sets. In total, 41 arthropod and mollusc sequences were removed from the initial set.

To build all phylogenetic trees, amino acid sequences were aligned using PASTA v1.8.6 (Mirarab et al., 2015). The alignments included 202 sequences previously described as aquaporins (Finn et al., 2014, 2015) to use as a backbone for identifying aquaporin subfamilies in our data set (Supp. Data 3). In addition, 3 previously identified malacoglyceroporins and malacoaquaporin (Kosicka et al., 2016) were also included when building aquaporin phylogenetic trees. All resultant alignments were trimmed using trimAL v1.4.1 (Capella-Gutiérrez, Silla-Martínez, & Gabaldón, 2009). We tested different trimming methods and gap thresholds to select the best one in terms of keeping fewer alignment positions to reduce computational time while keeping biologically informative sites (Fig S1). Final alignments were trimmed with a gap threshold of 0.01.

We created guiding trees for each of the alignments using the LG model in FastTree v2.1.11 (Price, Dehal, & Arkin, 2010), which were later used as starting trees for the ML inference of aquaporin trees with IQ-TREE v2.1.2 (Minh et al., 2020). We applied an LG protein matrix with a profile mixture model of 10 classes (C10), plus an additional class of empirical amino acid profile (F) and rate heterogeneity across sites using a discrete Gamma model with 4 categories (G). We also included ultrafast bootstrap (UFBoot) (Minh, Nguyen, & von Haeseler, 2013) and SH-aLRT test (Guindon et al., 2010) to obtain support values of resultant trees. The AqpM sequence of *Methanococcus maripaludis* (NCBI accession NP_988083), included in the backbone, was always used as an outgroup for tree rooting, as suggested in previous studies (Finn et al., 2014).

To avoid long branch attraction (LBA) due to their higher divergence and because our preliminary results using all aquaporins showed that they formed a monophyletic clade, entomoglyceroporins were analysed separately and excluded from the phylogeny that combined all animal Aqp1-like sequences.

We visualised the trees using iTOL (Letunic & Bork, 2021) and created the final figures with the R packages ggtree (Yu, Smith, Zhu, Guan, & Lam, 2017) and ggtreeExtra (Yu et al., 2021). A full list of packages used is included in the code available on GitHub (see Data availability section).

### Inference of a combined dated species tree

Time-scaled ultrametric trees were inferred independently for arthropods, molluscs and annelids by using a node-dating approach on species trees that followed the most recent topologies (Anderson & Lindgren, 2021; Ballesteros et al., 2022; Colgan, Ponder, Beacham, & Macaranas, 2007; Cunha & Giribet, 2019; Doğan, Schrödl, & Chen, 2020; Erséus et al.,2020; Fernández, Edgecombe, & Giribet, 2018; González et al., 2015; Johnson et al., 2018;Kallal et al., 2021; Kawahara et al., 2019; Kocot et al., 2011; Kocot, Halanych, & Krug, 2013;Kocot, Poustka, Stöger, Halanych, & Schrödl, 2020; Kocot et al., 2017; Kocot, Todt, Mikkelsen, & Halanych, 2019; Lee et al., 2019; Lindgren, 2010; Lozano-Fernandez et al.,2019; McKenna et al., 2019; Misof et al., 2014; Moles & Giribet, 2021; Ontano et al., 2021;Pabst & Kocot, 2018; Peters et al., 2017; Razkin et al., 2015; Richter, Richter, & Scholtz, 2001; Salvi & Mariottini, 2016; Schwentner, Richter, Rogers, & Giribet, 2018; Smith, 2021;Smith et al., 2011; Sun et al., 2020; Uribe, Colgan, Castro, Kano, & Zardoya, 2016; Uribe & Zardoya, 2017; Wang, Shi, Qiu, Che, & Lo, 2017; Wiegmann et al., 2011; Yang et al., 2018). Full details for the dating analysis of arthropods and molluscs are provided elsewhere (Aristide et al. in prep; Tonzo et al. in prep).

In the case of annelids, we dated the species tree using the Aqp1-like gene sequences in MCMCtree (Z. Yang, 2007). For that, we first inferred the annelid Aqp1-like gene tree using aquaporins annotated in (Mucciolo et al., 2021) and our inferred aquaporins for *E. andrei*, as our preliminary analyses showed that the aquaporin phylogeny is broadly concordant with the species tree topology at the level of aquaporin subclusters. Second, from this gene tree, we derived 16 sets of 1:1 orthologues with at least four species represented using PhyloPyPruner (Thálen, 2019), considering the phylogenetic interrelationships of annelid species as described in (Erséus et al., 2020). These orthologous sequences were then concatenated into a single matrix of 2,694 sites for their analysis and split into two partitions based on their evolutionary rate (measured as the sum of branch lengths divided by the number of tips in the respective gene tree). Prior distributions for node ages (see Supplementary Information in Github) were based on (Chen et al., 2020) for the annelid MRCA and in the divergence time intervals and fossil calibrations in (Erséus et al., 2020). Species *Romanchella perrieri* was removed by PhyloPyPruner and was therefore not included in the tree.

Vertebrate and onychophoran time trees were obtained from the literature (Baker, Buckman-Young, Costa, & Giribet, 2021; Irisarri et al., 2017) and pruned to keep only a selection of 13 vertebrate species whose aquaporins have previously been studied (Finn et al., 2014) and the 4 onychophorans (or closely-related species) that were included in MATEdb (Fernández et al., 2022), respectively.

Finally, the ultrametric trees of all 5 phyla were grafted into a manually-built backbone topology depicting the relationships among them using the RRphylo R package (Castiglione et al., 2018). Divergence times for the ancestral nodes in the backbone tree were obtained from (Dohrmann & Wörheide, 2017).

### Phylogenetic comparative methods to explore aquaporin evolution

In order to test whether there were differences in the mean total number of aquaporins between terrestrial and aquatic species, we implemented a phylogenetic generalised linear model using the phyloglm R package (Ho & Ané, 2014). Briefly, we fitted a phylogenetic Poisson regression model to our data describing the relationship between the number of aquaporins and habitat, accounting for the phylogenetic history of the species. For habitat, we classified species in two different ways: marine vs non-marine (i.e., freshwater, brackish and terrestrial), and terrestrial vs non-terrestrial (i.e., freshwater, brackish and marine). This classification allowed us to identify shifts in aquaporin number during marine to freshwater and terrestrial transitions, and marine and freshwater to terrestrial transitions. We tested for differences in each of the 4 main subtypes of aquaporins and in all aquaporins together.

## Results

### An expanded catalogue of aquaporins in arthropods, molluscs and onychophorans unveils the complex dynamics of aquaporin evolution

Our knowledge of animal aquaporins’ evolution is patchy. To close some gaps that affect our understanding of their potential involvement in terrestrialization, we first explored the aquaporin gene repertoire in arthropods, molluscs and onychophorans, since we had an extensive taxon sampling for each phylum. We identified a total of 2,303 arthropodan, 1,225 molluscan, and 52 onychophoran Aqps (Supp. Data 4) after annotating 310 arthropodan, 101 molluscan and 4 onychophoran genomic data sets. Only some Aqps from a few arthropod species and even fewer mollusc species had been previously annotated as aquaporins (Catalán-García et al., 2021; Colgan & Santos, 2018; Finn et al., 2015; Stavang et al., 2015). We also report here the first identification of Aqps in the phylum Onychophora.

As we wanted to include additional phyla to have a better understanding of the evolution of Aqp-coding genes in animals that have transitioned from the sea to the land, we searched the literature for additional aquaporin sequences in representatives from other phyla that underwent terrestrialization episodes (Nematoda, Tardigrada, Craniata and Annelida), obtaining a total of 4,230 aquaporins. Aquaporins of *C. elegans* (Huang et al.,2007) and *M. tardigradum* (Grohme et al., 2013) were classified into the four main subtypes by building a phylogenetic tree with the backbone sequences. For annelids and vertebrates, aquaporins had already been classified into the main groups, which were later confirmed in our phylogeny.

Our inferred phylogeny of all animal aquaporins (Fig S2) supports the four aquaporin clades previously defined in the literature (Abascal et al., 2014): aquaglyceroporins (Aqp3-like), classical aquaporins (Aqp1-like), aquaammoniaporins (Aqp8-like) and superaquaporins (Aqp11-like). Moreover, the loss of Aqp8-like in arthropods and Aqp1-like in *C. elegans* is recovered, as previously reported (Finn & Cerdà, 2015; Stavang et al., 2015), together with the identification of Aqp8-like loss in onychophorans, suggesting that the loss of Aqp8 may have occurred before the split between Onychophora and Arthropoda. Hexapod-specific glyceroporins (i.e. entomoglyceroporins or Eglps) clustered within arthropod Aqp1-like, showing their Prip-like origin as previously reported (Finn et al., 2015).

### Phylogenetic comparative methods unveil parallel aquaporin-coding gene duplications in non-marine animals and gene loss in terrestrial ones

The presence of Aqp clades specific to terrestrial animals, such as the Eglps in insects (Finn et al., 2015) or AQP2, AQP5 and AQP6 in tetrapods (Finn et al., 2014; Lorente-Martínez et al., 2018), suggests that the number of aquaporins found in terrestrial animals might be different than the number found in aquatic ones. To test whether there is a correlation between the number of aquaporins and the habitat in which extant species live, we obtained Aqp-coding genes identified in each species and performed a phylogenetic Poisson regression using the habitat as an independent variable. We tested two different hypotheses using this model. The first one is that the number of Aqp-coding genes in terrestrial species is different to that of aquatic ones, supporting that a copy number variation of Aqp-coding genes through gene gain, duplication or loss may have facilitated terrestrialization during aquatic-terrestrial transitions. The second hypothesis tested is that the number of Aqp-coding genes in marine species is different to that in non-marine species.

We first explored two independent tests with genomic data of the five phyla (Onychophora, Craniata (Vertebrata), Mollusca, Annelida and Arthropoda; five-phyla data set hereafter; Fig 2A). Tardigrades and nematodes were not included as we only analysed one species per phylum and lacked the representation we had in the others, which could bias the results. Moreover, since we had a dense taxon representation of most main lineages in arthropods and molluscs, we performed the tests independently for each of these two phyla as well. These tests were explored both taking into account all Aqp-coding genes and in each type of Aqp-coding genes independently, following the main four types described above (i.e., Aqp1-like, Aqp3-like, Aqp8-like and Aqp11-like).

**Figure 2.**
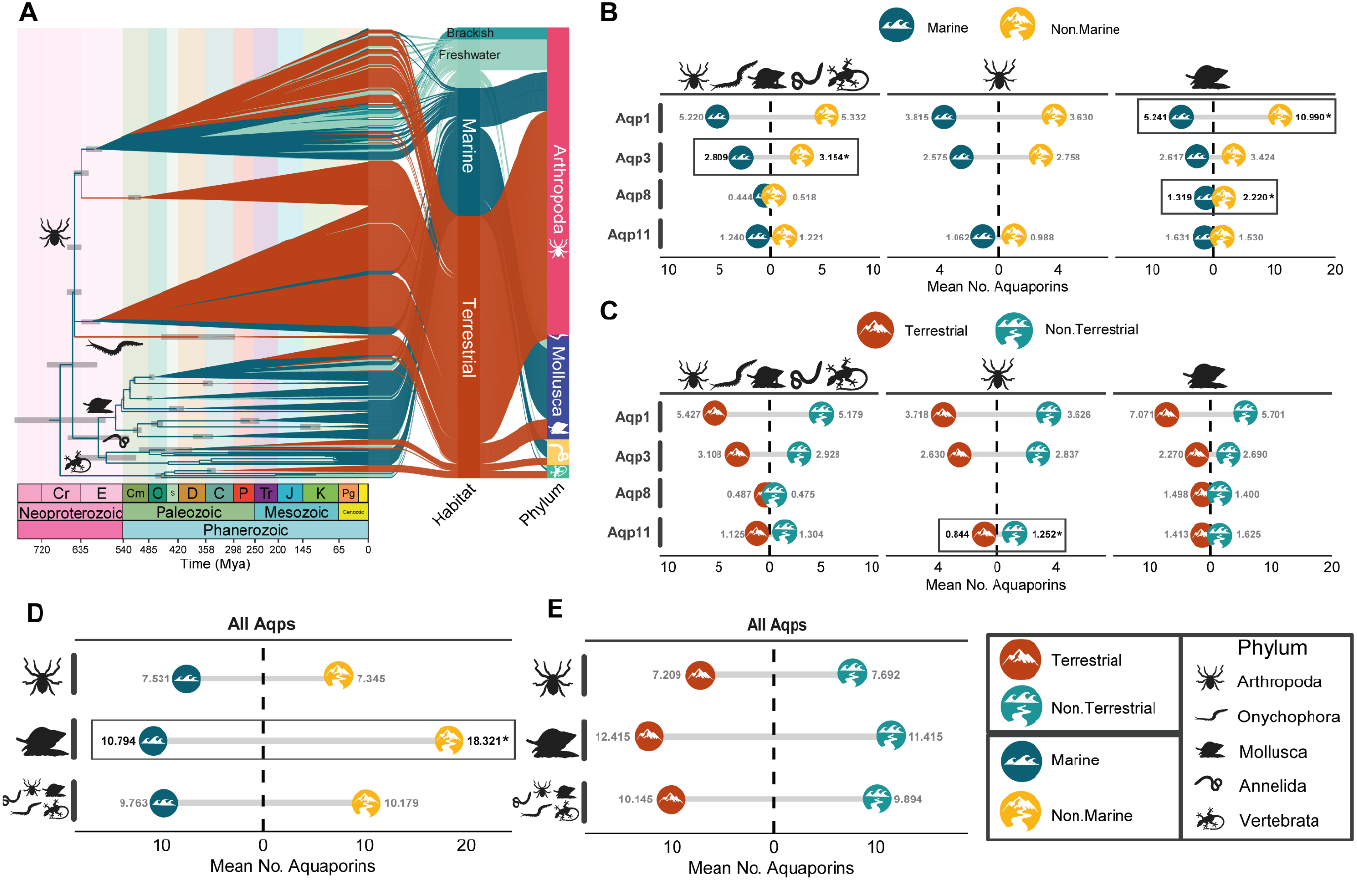
Phylogenetic comparative methods to understand aquaporin evolution. **A**. Ultrametric calibrated tree of the 483 species from 5 phyla included in the analyses. **B**. Phylogenetic Poisson regression results for marine (dark blue) vs non-marine (yellow) comparison in all animals, arthropods and molluscs in the four aquaporin subtypes (i.e. Aqp1-like, Aqp3-like, Aqp8-like and Aqp11-like). Significant results are highlighted in bold with an asterisk and inside a square. **C**. Phylogenetic Poisson regression results for terrestrial (dark orange) vs aquatic (blue) comparison in all animals, arthropods and molluscs in the four aquaporin subtypes (i.e. Aqp1-like, Aqp3-like, Aqp8-like and Aqp11-like). Significant results are highlighted in bold with an asterisk and inside a square. **D**. Phylogenetic Poisson regression results for marine (dark blue) vs non-marine (yellow) in all animals, arthropods and molluscs in all aquaporins. Significant results are highlighted in bold with an asterisk and inside a square. **E**. Phylogenetic Poisson regression results for terrestrial (dark orange) vs aquatic (blue) in all animals, arthropods and molluscs in all aquaporins.

Overall, our results showed statistical differences in the number of several aquaporin-coding gene subfamilies when we compared marine and non-marine species, pointing to a significantly higher number in non-marine species than in marine ones in most cases (Fig 2B and C). First, in the analyses of the five-phyla data set we observed that non-marine animals have a significantly higher number of Aqp3-like compared to marine species (Fig 2B, left). A new nomenclature for Aqp3-like subclades in the different animal phyla and the implications of its phylogenetic history in the context of animal terrestrialization is discussed in detail below (see also Table 1).

**Table 1.**
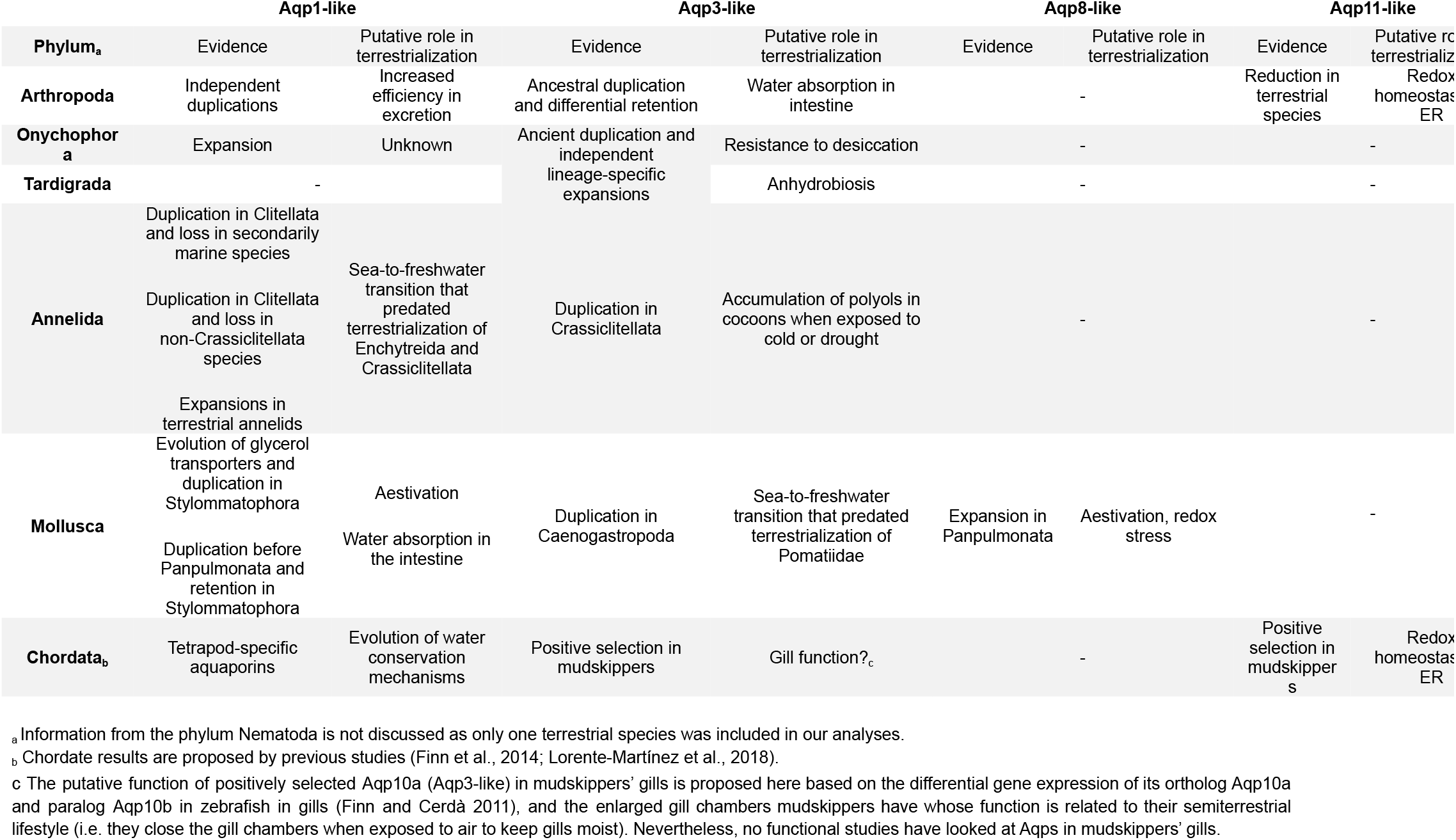
The role of aquaporin evolution in animal terrestrialization. Summary table of the obtained results and proposed hypotheses in this paper.

Regarding the phylum-specific analyses in molluscs, we recovered a significantly higher number of Aqp-coding genes in non-marine compared to marine species (Fig 2C). Independent analyses for each type of Aqp-coding gene revealed a significantly higher number in non-marine species of Aqp1-like and Aqp8-like (Fig 2B, right). A description of the phylogenetic relationships of Aqp1-like and Aqp8-like as well as their potential role in mollusc terrestrialization is discussed below.

In addition, our results indicated that terrestrial arthropods have a significantly lower mean number of Aqp11-like aquaporins than aquatic ones (Fig 2B, centre). Phylogenetic relationships among Aqp11-like aquaporins are represented in Fig S3, showing some lineage-specific duplications in secondarily aquatic species.

### Ancient independent duplications in Aqp-coding genes are generally correlated with the ‘out of the sea’ transition across animal phyla

#### a) Classical aquaporins (Aqp1-like)

The phylogeny of protostome classical aquaporins (Fig 3, Fig S4) reveals a complex pattern of gains and losses in the different aquaporin-coding gene clusters, with many originating early during animal evolution. We highlight two clades of Aqp1-like, which we coined as Aqp1-bila1 and Aqp1-bila2, and that include the majority of Aqp1-like animal sequences (except for Aqp1-ann1, Prip and Drip). Since deuterostomes (mostly vertebrates, Fig 3, in turquoise) cluster inside Aqp1-bila2, while being absent in Aqp1-bila1, we can infer that the duplication that gave rise to both Aqp1-bila clades occurred at the base of Bilateria, before the diversification into Deuterostomia and Protostomia. Both Aqp1-bila copies were further duplicated or lost independently across animal evolution. Aqp1-bila1 contains mainly genes from species within Spiralia, together with Onychophora and Tardigrada, while Aqp1-bila2 includes sequences from all animal phyla included in this study, except for nematodes, for which a lack of Aqp1-like sequences had previously been reported (Finn & Cerdà, 2015). It is worth mentioning that lineages that have the most profound adaptations to life on land (i.e. tetrapods and arthropods) underwent the loss of Aqp1-bila1 and only retain Aqp1-bila2.

**Figure 3.**
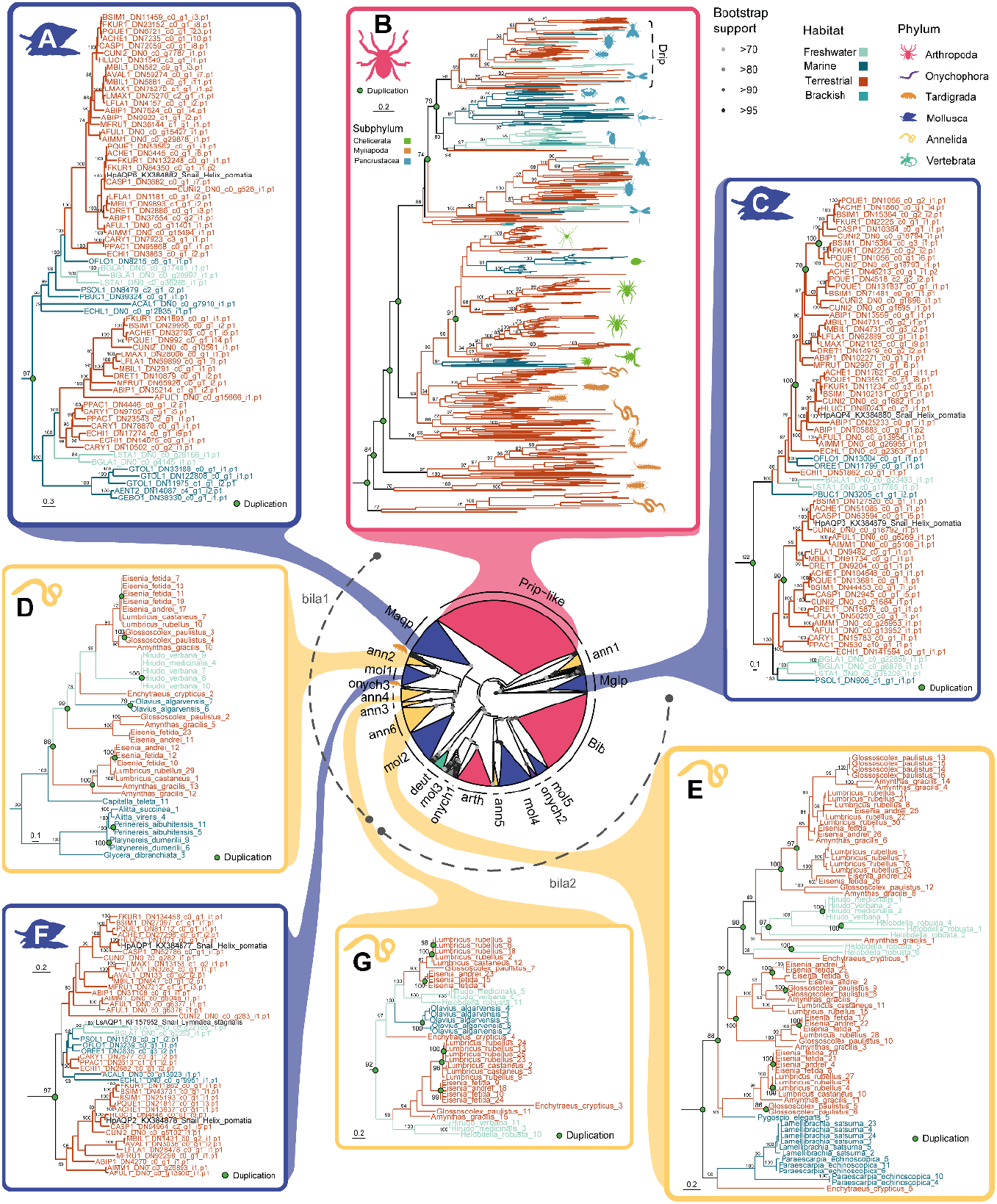
Consensus phylogenetic tree of 2,202 Aqp1-like plus 63 backbone sequences in 6 studied phyla. **A**. Both copies of malacoaquaporins (Maqp) in Panpulmonata. **B**. Prip-like and Drip clades in arthropods. **C**. Malacoglyceroporins’ duplication in Panpulmonata. **D**. Aqp1-ann2 expansion within Crassiclitellata. **E**. Aqp1-ann6 clade expansion in Crassiclitellata. **F**. Aqp1-mol1 duplication in Stylommatophora. **G**. Aqp1-ann4 subclade in Clitellata. Duplications are marked with a green dot. Bootstrap values are included only in nodes with support values higher than 70.

Classical aquaporins were also independently duplicated in terrestrial lineages (Fig 2B and C). Our data show a not fully resolved clade containing the insect-exclusive Drip and arthropod Prip-like clusters (Fig 3B). It appears that there were a series of expansions in the arthropod ancestor followed by a differential loss and independent duplications in the different arthropod lineages. The Prip-like pauropodan, symphylan and chilopodan myriapod classes retain two ancestral copies, while the origin of the second copy in diplopodan myriapods that retain more than one is more recent. Although previous studies have suggested that there has been an ancient whole-genome duplication (WGD) in Arachnopulmonata (Schwager et al., 2017), the Prip-like duplications that we recover do not suggest that as the origin of the Arachnopulmonata clades. On the contrary, the high clustering of all xiphosuran Prip-like duplicate sequences is indeed consistent with them being onhologs that originated after 3 rounds of WGD (Nong et al., 2021). Nevertheless, an independent phylogenetic tree we inferred using only arthropod sequences recovers a different topology, with hexapod Drip (including the previously-mentioned Drip insect-exclusive cluster and some additional early-splitting hexapod sequences) and Prip-like having emerged after the divergence of Pancrustacea from Myriapoda. However, this topology that suggests independent duplications in each of the three terrestrial arthropod groups is not supported by bootstrapping, and therefore these interrelationships should be taken with caution (Fig S5). In addition to Drip and Prip-like clades, it is noteworthy that the Aqp1-arth cluster (Fig 3) contains chelicerate and myriapod sequences, while it lacks pancrustacean sequences (except for some sequences from Malacostraca, Ostracoda, Cirripedia and a couple of earliest insect-splitting species - Fig S4).

Onychophorans, the only fully-terrestrial animal phyla, have three clusters of Aqp1-like sequences (Fig 3, in purple). One of them, Aqp1-onych2, harbours a gene duplication at the level of phylum, followed by several duplications and gene losses.

Regarding annelids, we observe 6 different clusters named Aqp1-ann1 to Aqp1-ann6 (Fig 3, in yellow). Among them are the Aqp1-ann6, Aqp1-ann4 and Aqp1-ann2 clusters (Fig 3E, 3G and 3D, respectively). Aqp1-ann6 include a wide expansion in terrestrial Crassiclitellata (i. e. earthworms), and loss of ancient copies in non-Crassiclitellata annelids, putatively related to their terrestrialization (Fig 3E). In addition, we observe a duplication of Aqp1-ann4 in Clitellata with a subsequent loss of one sub-copy in secondarily marine clitellates (*Olavius algarvensis*), as shown in Fig 3G. Regarding Aqp1-ann2 (Fig 3D), there is an expansion at the level of Clitellata with the subsequent loss of all copies but one in non-Crassiclitelata annelids.

In molluscs, we find 6 supported clades (Fig 3, in indigo), which we have named Aqp1-mol1-5, Maqp and Mglp (Malacoaquaporins and Malacoglyceroporins, respectively, in accordance with previous research on the stylommatophoran snail *Helix pomatia* (Kosicka et al., 2016). As mentioned before, gastropods are the only molluscs that have colonised land. When looking at the mollusc-specific classical aquaporin clusters, we find three groups of special relevance. The first one is a Stylommatophora-specific duplication within Aqp1-mol1 (Fig 3F). The origin of this duplication dates back to right before the origin of Panpulmonata (Tenctipleura), with one copy being lost in all non-Stylommatophoran lineages. (Kosicka et al., 2016) identified the two duplicates of this clade in *H. pomatia:* HpAqp1 and HpAqp2. Furthermore, we also report the expansion of Aqp1-mol2 in terrestrial Stylommatophora, freshwater panpulmonates, and the freshwater bivalves *Margaritifera margaritifera* and *Cristaria plicata* (Fig 3, Fig S4). Moreover, an additional duplication within Maqp (Fig 3A) occurred before the split of Gastropoda and Scaphopoda, with only one copy being retained in Scaphopoda, and two within panpulmonate gastropods. The rest of gastropods Maqp are not clustered within this clade.

#### b) Aquaamnioporins (Aqp8-like)

We identified several lineage-specific duplications of Aqp8-like in *C. elegans*, annelids and molluscs (Fig S6). In the case of annelids both terrestrial and aquatic species show duplications within this aquaporin subtype. However, not all molluscan lineages showed duplications. For instance, gastropods contain only one copy of Aqp8-like, except for the terrestrial gastropod *Pomatias elegans*. In addition, while marine bivalves contain only 1 or 2 copies, freshwater bivalves contain 3 or even 4. These two examples point to the putative role of these expansions in their transition from the marine environment.

#### c) Aquaglycerolporins (Aqp3-like)

Compared to the phylogeny of Aqp1-like (Fig 3), where deuterostome aquaporins were nested within protostome aquaporins showing a shared origin, deuterostome and protostome Aqp3-like form two separate clades with independent evolutionary histories (Fig 4, Fig S7). Despite using a different methodology, our results of vertebrate Aqp3-like are mostly consistent with previous studies (Finn et al., 2014), except for the recovery of AQP10 and AQP7 as sister clades. Within protostomes, there is a further split into two clades, which we named Aqp3-prot1 and Aqp3-prot2. Aqp3-prot1 includes representatives of all invertebrate phyla included in this study. On the contrary, Aqp3-prot2 is mostly composed of arthropod sequences but includes as well some sequences from some mollusc and nematode species.

**Figure 4.**
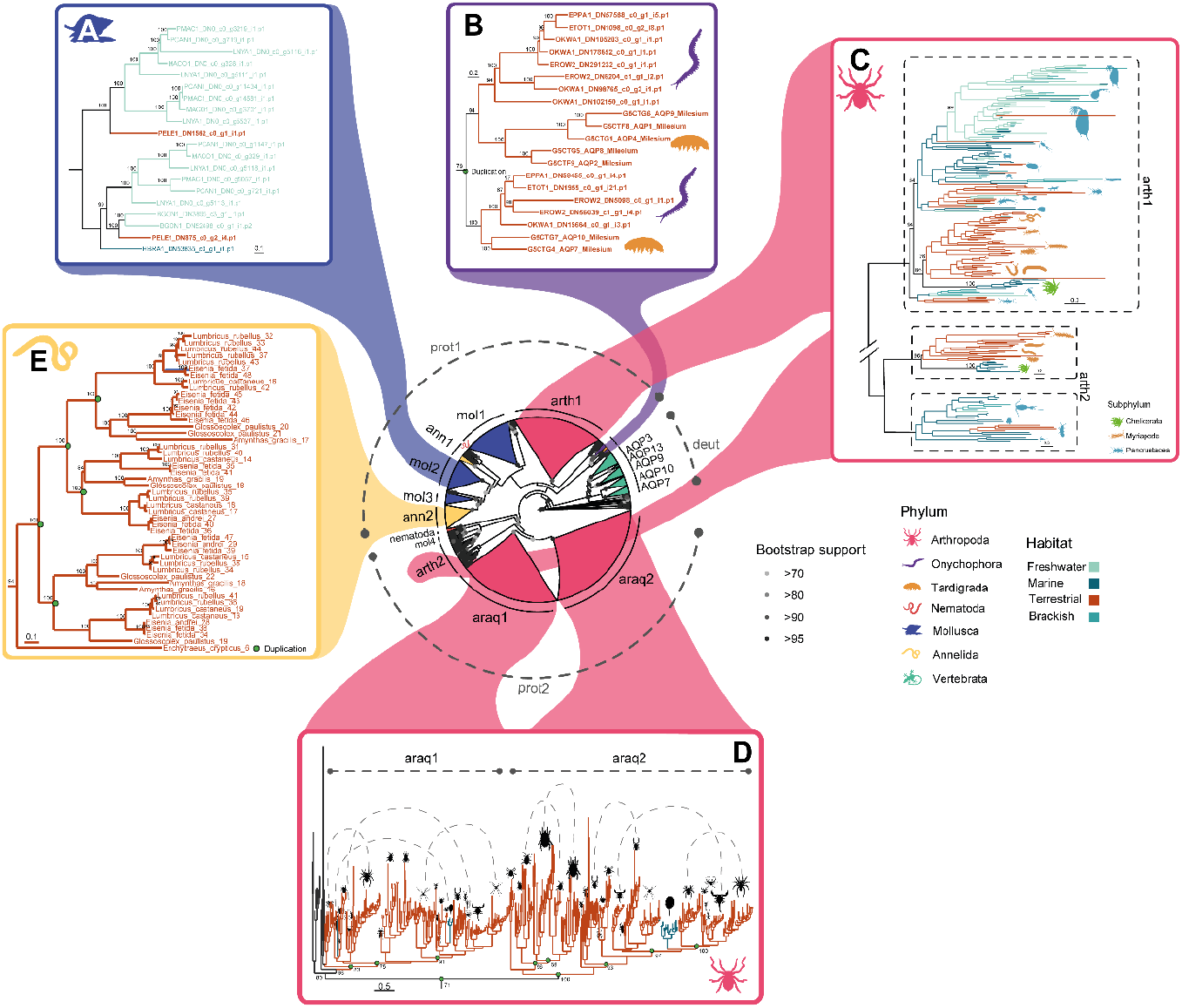
Consensus phylogenetic tree of 1,108 Aqp3-like plus 68 backbone sequences in the 7 studied animal phyla. **A**. Aqp3-mol2 clade in Caenogastropoda shows a duplication in this freshwater-origin subclass. **B**. Aqp3-like paralogs in Onychophora and Tardigrada. **C**. Aqp3-arth1 and Aqp3-arth2. **D**. Aqp3-araq1 and Aqp3-araq2 arachnid paralogs. **E**. Aqp3-ann2 expansions within Crassiclitellata. Duplications are marked with a green dot. Bootstrap values are included only in nodes with support values higher than 70.

Zooming in the phylogenetic tree inferred for Aqp3-like, we recovered several ancient duplications in the nodes where a habitat transition occurred. Previous studies have reported independent duplications of Aqp3-like within chelicerates (Finn et al., 2015), myriapods and copepods (Catalán-García et al., 2021). However, their results could have been biassed due to the omission of other major arthropod lineages in their inferences. Here, we overcome this limitation by having representatives of virtually all arthropod orders analysed together. Our results pointed to an ancestral duplication that was independently retained in some groups and lost in others (Fig 4 and Fig 4C), instead of inferring independent within-lineage duplications. For instance, (Finn et al., 2015) report that two clades of Aqp3-like evolved in chelicerates based on their phylogeny with only spider, scorpion and acari sequences. When including the rest of the arachnid orders, such as xiphosurans and pycnogonids, results show that the duplication they reported is arachnid-specific (with xiphosurans nested within arachnids, Fig 4D) and derives from one of the two ancient copies of Aqp3-like that are present in arthropods, while having lost the other one. On the contrary, Pycnogonida, the earliest-splitting chelicerate lineage, has retained both copies. This is consistent with a single terrestrialization event in arachnids (considering a secondary later colonisation of water by xiphosurans as discussed in (Ballesteros et al., 2022)), where the duplication of Aqp3-like may have helped chelicerates to transition to a different niche. These Aqp3-like copies subsequently underwent independent duplications in both clades, with two subclades in Xiphosura, Ricinulei, Opiliones and Arachnopulmonata (which we named Aqp3-araq1, not likely the result of their WGD; Fig. 4D), and two subclades in Acariformes, Scorpiones, Pseudoscorpiones and Araneae (which we named Aqp3-araq2, Fig 4D). Regarding myriapods, (Catalán-García et al., 2021) reported that Aqp3-like genes within Pauropoda, Symphyla and Diplopoda clustered in two separate clades, the ones that we inferred in this study as ancient duplications. Symphyla and Diplopoda retain both ancient copies (named Aqp3-arth1 and Aqp3-arth2 in Fig 4C), while Chilopoda and Pauropoda only retain Aqp3-arth1. Last, but not least, we reproduce the pancrustacean results obtained by (Catalán-García et al., 2021) regarding the presence of two copepod Aqp3-like clusters, one of them as a sister group to Thecostraca and Arguloida Aqp3-like (in which we also report a close relationship to Collembola), and the other clustered within the rest of Pancrustacea (Fig 4C). We also obtain an early-branching clade of Aqp3-arth1 composed of freshwater copepods and terrestrial diplurans, albeit recovered with low bootstrap support. Noteworthy, we also confirm that holometabolous insects have lost Aqp3-like sequences in favour of a potentially more efficient insect-specific Aqp1-like-derived glycerol transporter, as previously reported by (Finn et al., 2015). Moreover, we observe a Branchiopoda-specific duplication in Aqp3-arth1 (Fig 4C) that was also reported in previous studies (Catalán-García et al., 2021;Finn et al., 2015; Stavang et al., 2015).

Aqp3-like sequences from the other two members of Panarthropoda (Onychophora and Tardigrada) form a separate cluster from Arthropoda, with an intertwined evolutionary history. Fig 4B shows a shared duplication between Onychophora and Tardigrada, which suggests independent gene loss of both copies in arthropods. Each of these subclades underwent subsequent duplications, with an expansion within the onychophoran family Peripatopsidae. However, if the tardigrade copies have a species-specific or a much older origin cannot be pinpointed with the current taxonomic representation.

In the case of terrestrial annelids (Clitellata), our results show highly supported gene duplications of Aqp3-ann2 in earthworms (Crassiclitellata), but not in the terrestrial enchytraeids (Fig 4E). It is noteworthy that the exclusively terrestrial Aqp3-ann2 is clustered together with Aqp3-mol3, which contains aquaporins from mostly terrestrial panpulmonate gastropods, Caudofoveata and Cephalopoda.

Relative to molluscs, we report a duplication of Aqp3-mol2 in the freshwater-origin molluscan subclass Caenogastropoda (Fig 4A), that was subsequently lost in the only marine species we included in our data set (*Hinea brasiliana*). The presence or absence of this duplication in other Caenogastropoda marine species remains to be studied, but the presence of both copies in the terrestrial *P. elegans* suggests that it was at least retained in the common ancestors of this species with other marine members of the group. Interestingly, contrary to Stylommatophora, *P. elegans* has not lost Aqp3-mol1.

### Convergent evolution of Aqp1-like glycerol transporters in Panpulmonata and Insecta via gene duplication

In addition to Aqp3-like, previous studies have proven the ability of some terrestrial-specific Aqp1-like proteins to transport glycerol. Such is the case of entomoglyceroporins, insect-specific Aqp1-like derived glycerol transporters (Finn et al.,2015), and AQP6, a tetrapod-specific classical aquaporin that can also transport glycerol (Holm, Klaerke, & Zeuthen, 2004). Moreover, the rise of entomoglyceroporins (Eglps) was accompanied by the loss of Aqp3-like in holometabolous insects. Here, we describe a similar pattern in molluscs: the rise of Aqp1-like malacoglyceroporins (Mglp, Fig 3C), and the lack of panpulmonate representatives in Aqp3-mol1 and Aqp3-mol2 (Fig 4). Mglps are glycerol transporter aquaporins which have been previously described as mollusc-specific by (Kosicka et al., 2016). Notwithstanding, our larger taxonomic representation of molluscs restricts its specificity to gastropods, more specifically to Euthyneura. In addition, the clade contains a duplication at its root, right before the origin of Panpulmonata (Euthyneura), gastropods that have mantle-derived lungs. Non-marine Panpulmonata are in fact the only ones that retain both copies after the duplication, suggesting a potential association with their transition out of the sea.

Regarding Eglps, we observe that the most diverse holometabolous insects (e.g., Diptera, Coleoptera, Lepidoptera and Hymenoptera) contain two or more groups of entomoglyceroporins (Fig S8), compared to non-holometabolous insects. Further investigations including a denser taxonomic representation of insects will help shed light on this observation and pinpoint the exact origin of each duplication. Moreover, some species or groups of insects that colonised freshwater or the sea contain additional copies of Eglps (i.e. *Cloeon dipterum, Gerris buenoi* and *Anopheles gambiae* species and the Ephrydidae dipteran family). The secondarily marine *Anopheles merus* lacks Eglps, which may be the result of a secondary loss.

## Discussion

### Potential implications of parallel gene duplication and loss of aquaporin coding genes for animal terrestrialization

The transition of life from the sea to the land, either directly or through a freshwater route, involved the acquisition of physiological osmoregulatory adaptations to avoid water loss and keep the osmotic homeostasis of internal fluids. Non-marine species are osmoregulators compared to marine species, which are osmoconformers (except for teleosts due to their hypothetical freshwater origin (Carrete Vega & Wiens, 2012; Rankin & Davenport, 1981). In addition, the molecular function of aquaporins as a transporter of water and other molecules has been previously associated with osmoregulation (Cabrero et al.,2020; Finn et al., 2014; Spring et al., 2009; Wittekindt & Dietl, 2019; Yi et al., 2011) and even with terrestrialization in tetrapods (Finn et al., 2014). Here, we explored the Aqp-coding genes gain, duplication and loss across several animal phyla that conquered land, providing evidence of ancient parallel duplications at the base of the lineages that transitioned out of the sea. As the aquaporin transporting properties have only been sparsely explored in these non-tetrapod lineages through functional experiments, we hypothesise the underlying role aquaporin duplications may have had on the emergence of osmoregulation during animal terrestrialization based on this experimental evidence. Notwithstanding, we acknowledge that these interpretations should be understood as phylogenomically-informed hypotheses and not as scientific claims, and that further studies are necessary to test them in depth.

The role of polyols (*e.g*. glycerol) and sugars (*e.g*. trehalose) as osmolytes in animal osmoregulation has been known for years (Christoph, Beck, & Neuhofer, 2007; Lamitina, Morrison, Moeckel, & Strange, 2004; Thorat, Gaikwad, & Nath, 2012), as well as the ability of Aqp3-like to transport these molecules under osmotic stress(Gil, Jarius, von Kries, & Rohlfing, 2017; Laforenza, Scaffino, & Gastaldi, 2013). Thus, during animal paths out of the sea towards land or freshwater habitats, a higher number of Aqp3-like may have provided a more efficient osmolyte flux in fluids to counteract water loss (Fig 2B, left). Further studies on the transporting properties of Aqp3-like in marine and non-marine species will help to elucidate this.

Arthropods are the most successful animal phylum to colonise land, with 3 independent ancient ‘out-of-sea’ transitions (Dunlop, Scholtz & Selden, 2013). Our comparative phylogenetic analysis reported a significantly lower number of Aqp11-like in terrestrial arthropods (Fig 2B, centre). Based on the few experimental evidence of Aqp11-like supporting the role of this intracellular aquaporin in keeping redox/pH homeostasis in the endoplasmic reticulum (ER) (Bestetti et al., 2020; Ishibashi, Tanaka, & Morishita, 2021;Nozaki, Ishii, & Ishibashi, 2008), we can hypothesise a potential link of gene loss in Aqp11-like coding genes with a more efficient homeostasis in land arthropods. During hypoxia, Reactive Oxygen Species (ROS), such as H_2_O_2_, are generated in the ER. As the levels of oxygen are higher on land, the transition from the sea may have removed the need for H_2_O_2_ transport through Aqp11-like due to a lower production of ROS, which may have triggered gene loss in Aqp11-like coding genes. This hypothesis is further reinforced by the identification of several independent duplications of Aqp11-like coding genes in aquatic arthropod species - i.e. xiphosurans and mayflies (Fig S3), all of them occurring in species that had habitat transitions back to aquatic habitats (from land to sea, and land to freshwater, respectively). In addition, we detect Aqp11-like duplications in freshwater copepods and crayfish, which may be related to their transition from sea to freshwater. Further functional studies are most needed to understand the functional role of this enigmatic type of aquaporins and their role during arthropod terrestrialization.

Duplications and loss of other aquaporins may have also helped arthropods to successfully conquer land. We report here an ancient duplication of Aqp1-like with a secondary loss of Aqp1-arth in most pancrustacea (especially in hexapods, Fig 3). This loss could be a consequence of the origin of the hexapod-exclusive Drip cluster, as functional studies in *D. melanogaster* have demonstrated the high efficiency of water transport in Malpighian tubule cells expressing Prip and Drip, as well as their importance in desiccation resistance (Cabrero et al., 2020). The functional efficiency of Prip-like duplicates in the other land-dwelling arthropod lineages (arachnids and myriapods) remains to be assessed. Furthermore, our phylogenetic analysis of Aqp3-like shows an ancient gene duplication at the root of arachnids (Fig 4D). We could hypothesise that this new copy might have replaced the ancient Aqp3-like if it were neo- or subfunctionalized into a more efficient transporter, as reported with other aquaporins (Finn et al., 2015). Nevertheless, the subfunctionalization of *Lepeophtheirus salmonis’* Aqp3-like in the enterocytes reported by (Catalán-García et al.,2021) may suggest a similar role during intestine water absorption in other arthropods. Lastly, the presence of a Aqp3-like duplication in Branchiopoda (Fig 4C), crustaceans of marine origin that colonised brackish and freshwater environments (Oakley, Wolfe, Lindgren, & Zaharoff, 2013), with Thecostraca having lost one of the copies, reinforces our hypothesis of the role of aquaporins in osmoregulation.

The two phyla most closely related to arthropods, onychophorans and tardigrades, both of them having colonised land independently, also show Aqp-coding gene duplications in Aqp1-like (Fig 3) and Aqp3-like (Fig 4B). However, only the role of *M. tardigradum* Aqp4 (Aqp3-like) in anhydrobiosis has been previously put forward(Schokraie et al., 2012), hinting at the putative role onychophoran Aqp3-like may have in desiccation resistance and, therefore, in non-arthropod Panarthropoda terrestrialization. The role of Aqp1-like duplications in onychophoran physiology can only be hypothesised based on comparisons with other animals in the absence of functional studies.

The transition from a marine to a freshwater environment in clitelates (Erséus et al.,2020) may have been accompanied by the reshaping of their osmoregulatory capabilities. Here we report a wide expansion of Aqp1-ann6 in terrestrial clitelates, and loss in non-Crassiclitellata annelids (Fig 3E). Clitellata class is characterised by the presence of a clitellum that secretes a cocoon. Cocoons are egg sacs that nourish the developing embryos, whose structure and composition differ between clitellates. Previous studies have shown that earthworm but not enchytraeid cocoons accumulate polyols when exposed to cold or drought (Bauer, Worland, & Block, 2001; Holmstrup, 1995). Thus, successive highly supported gene duplications of putative polyol transporters Aqp3-ann2 in Crassiclitellata, but not enchytraeids (Fig 4E), suggest their putative role in this cocoon’s physiological response to drought and cold. Resistance to drought and cold might have facilitated their transition to the heterogeneous environmental conditions of land. Nevertheless, in the lack of functional studies of these aquaporins or the organs in which they are expressed, we can only suggest putative roles in annelid terrestrialization based on the osmoregulatory role of aquaporins in non-annelid organisms (Cabrero et al., 2020; Catalán-García et al., 2021; Finn et al., 2014;Gil et al., 2017; Laforenza et al., 2013; Zieger et al., 2022).

The transition of gastropods to land, either through a freshwater path or marginal environments (Krug et al., 2022; van Straalen, 2021; Vermeij & Watson-Zink, 2022), could have been facilitated by aquaporins evolution. Our independent comparative phylogenetic analyses for each type of Aqp-coding gene revealed a significantly higher number in non-marine species of Aqp1-like and Aqp8-like (Fig 2B, right). As the number of aquaporins in terrestrial species compared to non-terrestrial ones is not significant, this suggests that, as in the case of aquaglyceroporins in the five-phyla analysis, aquaporin-coding gene duplication potentially enabled to cope with osmotic stress during the transition of molluscs to freshwater and land. In fact, this potential association is further reinforced by the identification of Aqp1-like, Aqp3-like and Aqp8-like duplications (Figs 3A, C and F, 4A, and S6, respectively), and previous functional experiments. To give an example, we recovered the expansion of Aqp1-mol2 in freshwater bivalves (Fig S4) as sister group to a previously reported expansion in *Dreissena rostriformis* (Calcino et al., 2019), which was suggested to have a role in the transition from sea to freshwater as well as having a critical role during *D. rostriformis’* early cleavage (Zieger et al., 2022).

Regarding terrestrial gastropods, Stylommatophora-specific duplication within Aqp1-mol1 that contain previously described HpAqp1 and HpAqp2 (Kosicka et al., 2016), may have contributed to the success of this order in its transition to land via sub- or neofunctionalization to maximise water reabsorption in the intestine. This hypothesis assumes that *HpAqp2* and *HpAqp1* orthologues preserve the same function in the rest of stylommatophoran species studied here, where *HpAqp2* orthologues would be highly expressed in the intestine, an organ whose main function is water reabsorption (Forester,1977), but *HpAqp1* lowly expressed in foot, kidney and intestine (Kosicka et al., 2016). In addition, one of the Maqp (Fig 3A) and one of the Mglp (Fig 3c) paralogs contain the aestivation-related *HpAqp6* and *HpAqp3*, respectively, which are expressed in the foot during snail aestivation (Kosicka et al., 2020), one of the most relevant mechanisms of adaptation to humidity stress (Schweizer, Triebskorn, & Köhler, 2019). This functional relationship to the drought-related behaviour evokes the putative role this pulmonate-specific duplication might have had on the panpulmonate terrestrialization events.

Lastly, in contrast to Aqp1-like or Aqp3-like, not much is known about Aqp8-like, except for their ability to transport water and ammonia and their expression in the inner membrane of mammalian mitochondria (Calamita et al., 2005; Lee & Thévenod, 2006; Soria et al., 2010). Its relevant contribution to mitochondrial respiration (Ikaga et al., 2015) implies its importance for normal cell functioning and stress tolerance (Sokolova, 2018). Thus, the duplication of Aqp8-like in *C. elegans*, freshwater bivalves and the terrestrial gastropod *P. elegans* (Fig S6) suggests their importance during the ‘out of the sea’ transition. Further studies that include more nematodes, as well as functional characterisation of this aquaporin will shed light into its role during this major evolutionary transition.

## Conclusions

The molecular function of aquaporins as transporters of water and other molecules makes them an ideal system to interrogate the evolution of osmoregulation during animal terrestrialization. Here, we provide phylogenetically-informed evolutionary evidence of gene duplication and loss in Aqp-coding genes and discuss their potential key role during the evolution of terrestrial animal life. By comparing our phylogenetic results with previously described functional evidence, we hypothesise in which physiological processes aquaporins might have contributed to the transition of the different animal phyla from the sea to freshwater or land, summarised in Table 1. Overall, our results suggest that ancient independent expansions of aquaporins in non-marine animals may have played a key role in the adaptation to life on land through the evolution of more sophisticated osmoregulatory mechanisms, such as a potential role of Aqp1-like in molluscan aestivation or the putative role of Aqp3-like in earthworm cocoon resistance to drought. Our results also show the complexity of aquaporin evolution in animals, highlighting the importance of taxon sampling when studying ancient gene families. All in all, our study sets the road towards a deeper understanding of one of the many pieces in the evolutionary puzzle of animal terrestrialization: the emergence and reshaping of osmoregulation.

## Data availability statement

All supplementary information and code are included in the Github repository: https://github.com/MetazoaPhylogenomicsLab/Martinez_Redondo_et_al_2022_aquaporins

## Author contribution

GIMR and RF conceived the study. GIMR performed the analyses and designed the figures. GIMR and CSG annotated aquaporins. PBG and VT assisted aquaporin annotation. LA designed the Poisson regression analysis. LA and VT inferred the ultrametric trees. LA and GIMR performed the Poisson regression analysis. GIMR and RF interpreted the results. GIMR and RF wrote the first version of the manuscript. RF provided resources and supervised the study. All authors revised and approved the final version of the manuscript.

## Acknowledgements

GIMR acknowledges the support of Secretaria d’Universitats i Recerca del Departament d’Empresa i Coneixement de la Generalitat de Catalunya and European Social Fund (ESF) ‘Investing in your future’ (grant 2021 FI_B 00476) and the CSIC 2020 JAE Intro programme (JAEINT_20_02210). LA acknowledges funding from a Juan de la Cierva-Formación (grant agreement FJC2019-042184-I funded by MCIN/AEI/10.13039/501100011033). PBG was supported by an FPI grant (grant agreement no. BES-2017-081050) financed by MCIN/AEI /10.13039/501100011033 and by ESF ‘Investing in your Future’. RF acknowledges support from the following sources of funding: Ramón y Cajal fellowship (grant agreement no. RYC2017-22492 funded by MCIN/AEI /10.13039/501100011033 and ESF ‘Investing in your future’), Agencia Estatal de Investigación (project PID2019-108824GA-I00 funded by MCIN/AEI/10.13039/501100011033) and the European Research Council (this project has received funding from the European Research Council (ERC) under the European’s Union’s Horizon 2020 research and innovation programme (grant agreement no. 948281)). We also thank Centro de Supercomputación de Galicia (CESGA) for access to computer resources, and particularly Pablo Rey for his assistance and guidance. We thank Julien Clavel for helping us with the poisson regression analyses.

## Supplementary figures

**Figure S1.**
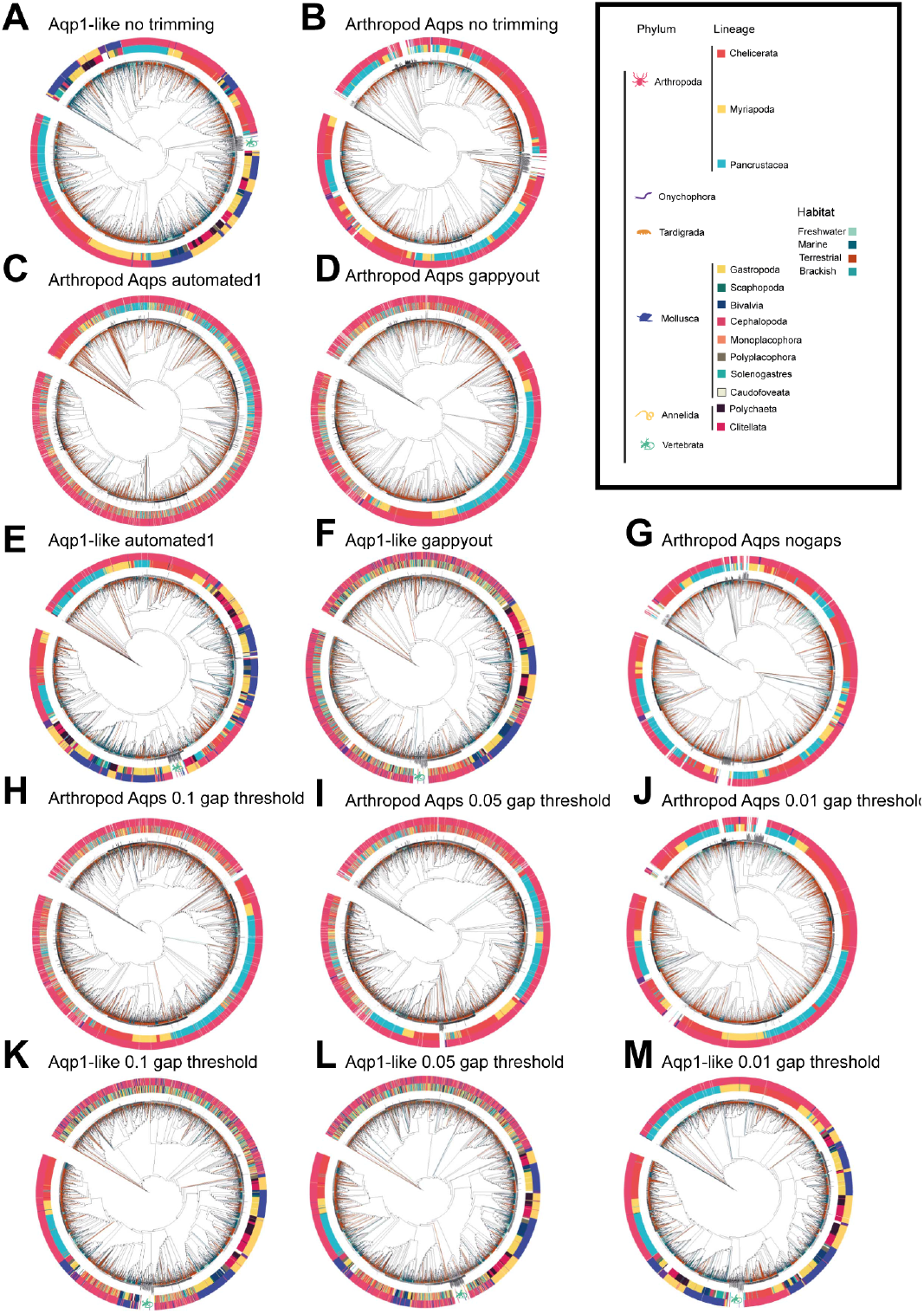
ML phylogenies built for all Aqp1-like and all arthropod Aqps to test the effect of alignment trimming using trimAL. Outer circumference represents the phylum and inner circumference represents the lineage inside each phylum. Different trimming options affect the formation of Aqp clades, with more extensive trimming options generating clades formed by Aqps of mixed origins. **A,B**. Without applying any trimming step. **C,E**. trimAL automated1 option, which opted for the strict option in both cases based on sequence number and identity score. **D,F**. trimAL gappyout option, which automatically selects a gap threshold (-gt) based on gap distribution. **G**. Removing all sequences that have a gap in arthropod Aqps. Applying this option to Aqp1-like alignment produced the same alignment as the one without any type of trimming. **H-M**. Selecting 3 different gap thresholds (1 - fraction of sequences with a gap allowed).

**Figure S2.**
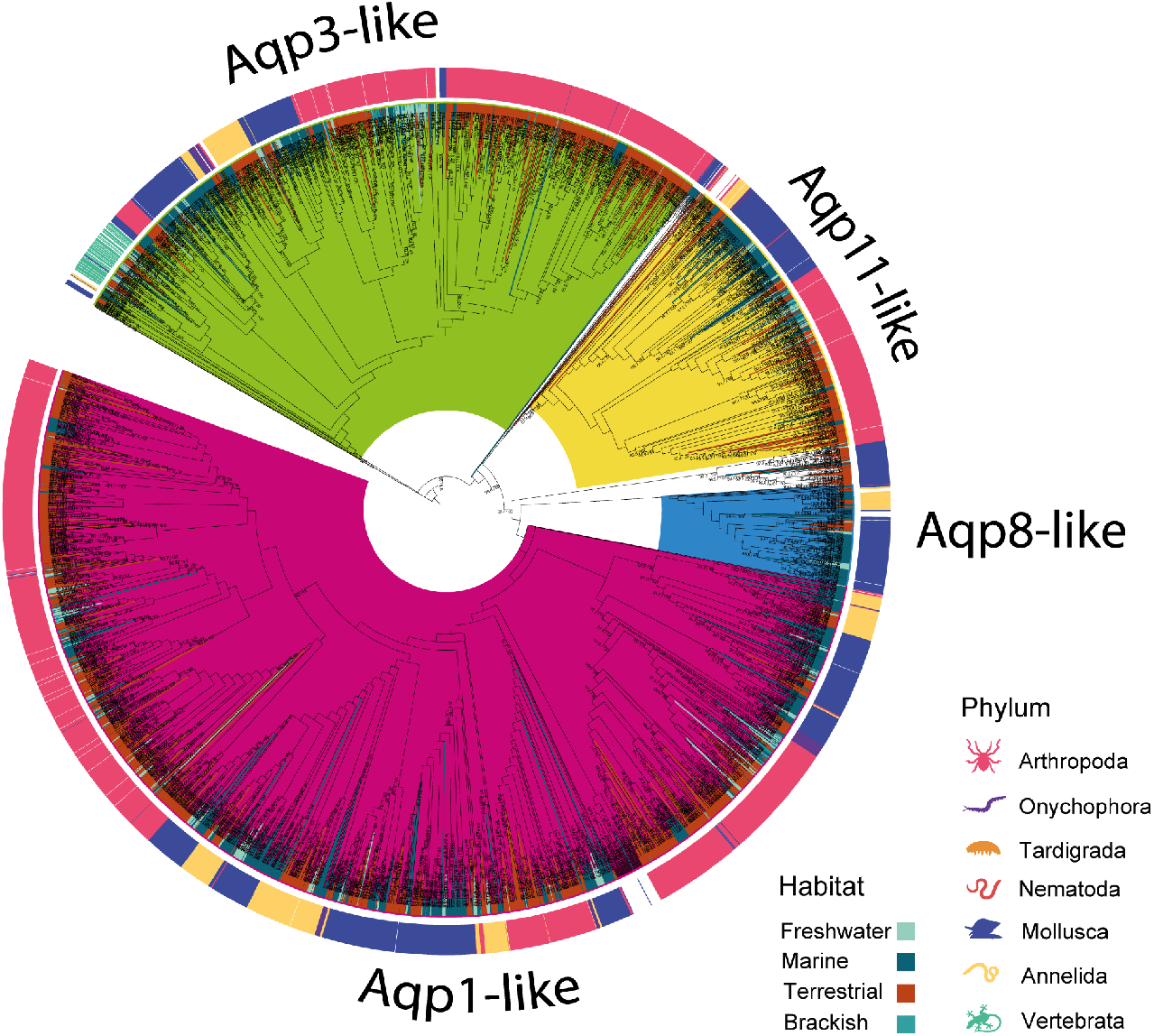
Consensus phylogenetic tree of all Aqps in the 7 studied animal phyla.

**Figure S3.**
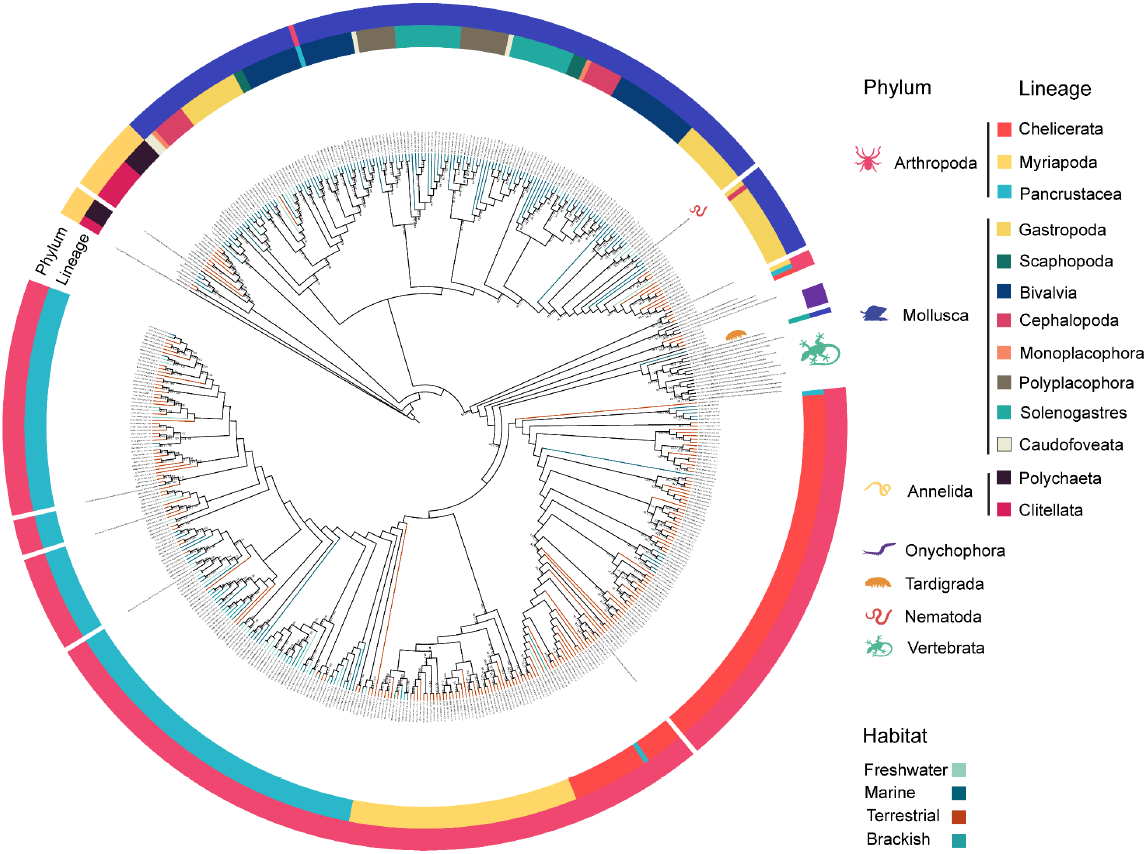
Phylogenetic tree of 481 Aqp11-like plus 27 backbone sequences in the 7 studied animal phyla.

**Figure S4.**
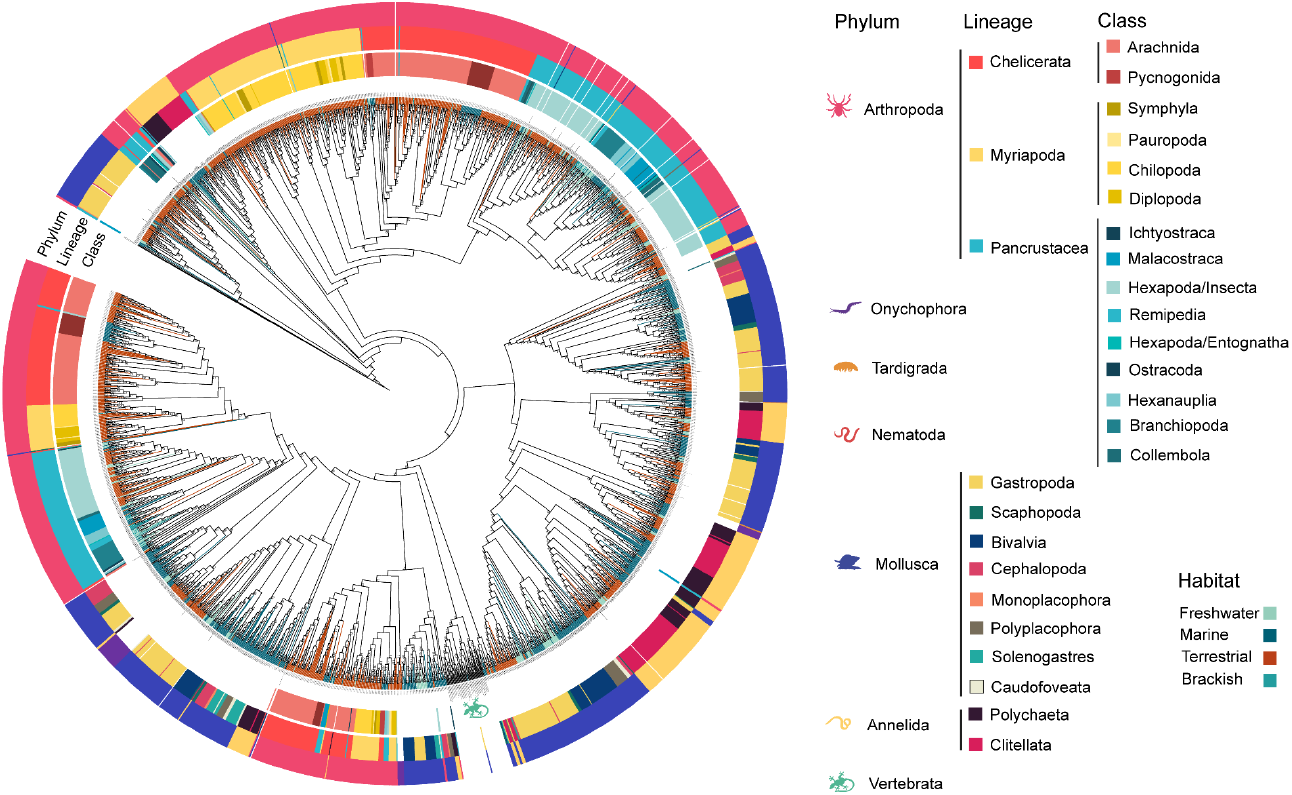
Uncollapsed consensus phylogenetic tree of 2,202 Aqp1-like plus 63 backbone sequences in 6 studied phyla.

**Figure S5.**
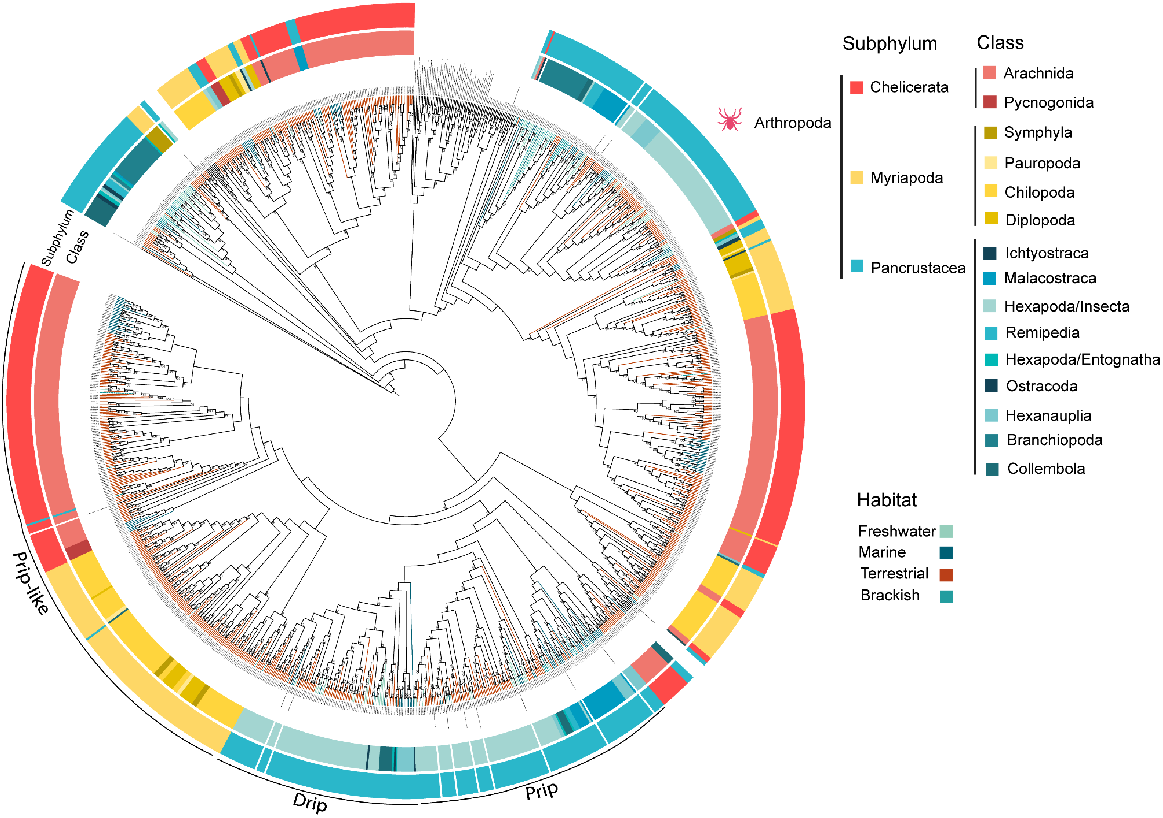
Consensus phylogenetic tree of 1,156 arthropod Aqp1-like plus 64 backbone sequences.

**Figure S6.**
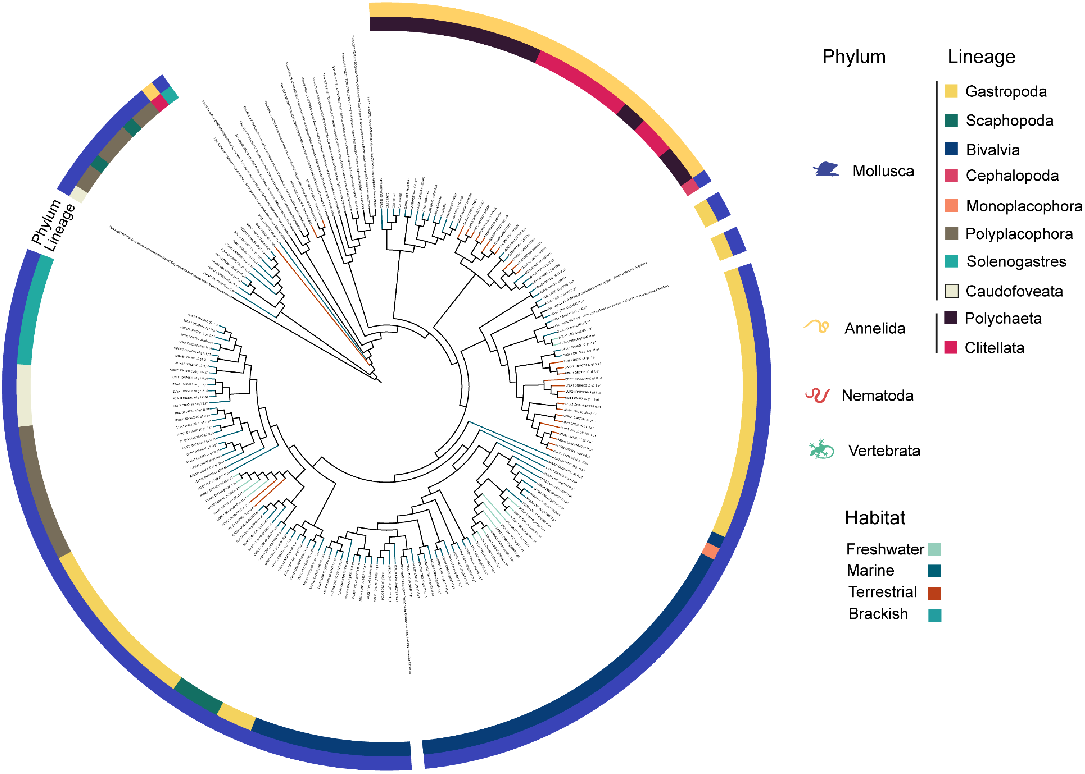
Phylogenetic tree of 162 Aqp8-like plus 19 backbone sequences in 4 studied animal phyla.

**Figure S7.**
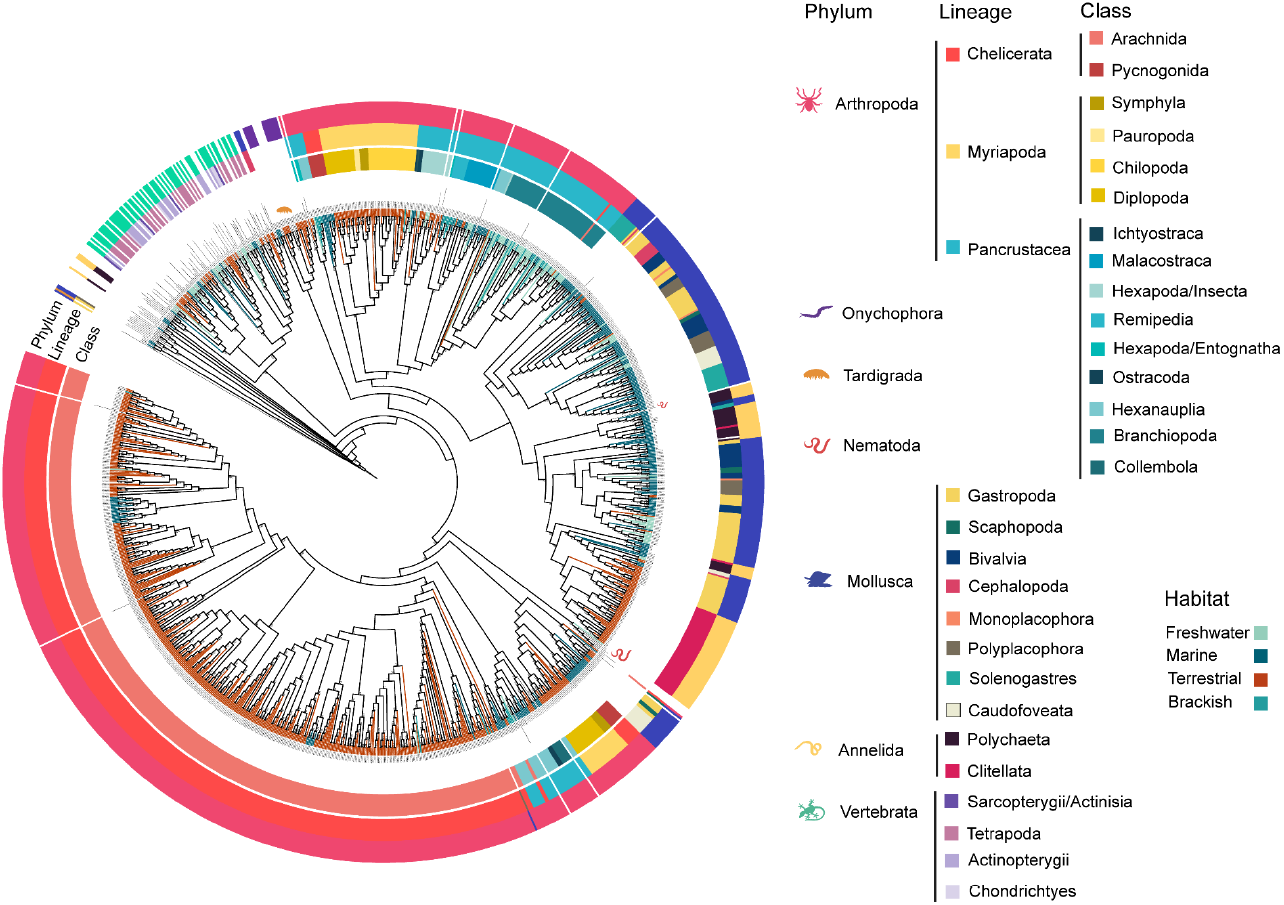
Uncollapsed consensus phylogenetic tree of 1,108 Aqp3-like plus 68 backbone sequences in the 7 studied animal phyla.

**Figure S8.**
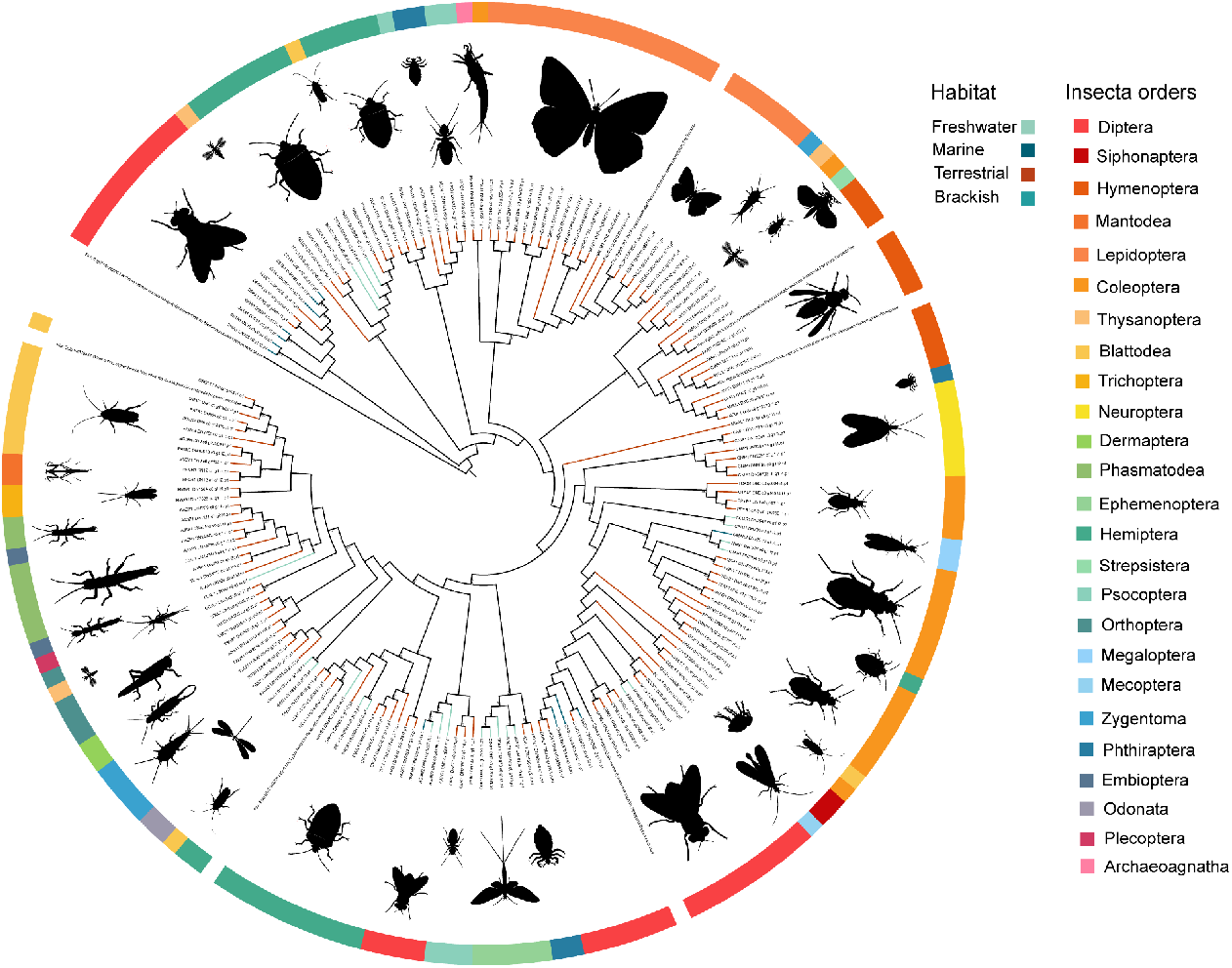
Phylogenetic tree of 176 Eglps plus 7 backbone sequences in the insects included.

