## Supplementary material for "Parallel duplication and loss of aquaporin-coding genes during the ‘out of the sea’ transition as potential key drivers of animal terrestrialization": Suppl. Figure 1

### Supplementary figures

**Figure S1. Consensus phylogenetic tree of all Aqps in the 7 studied animal phyla.**

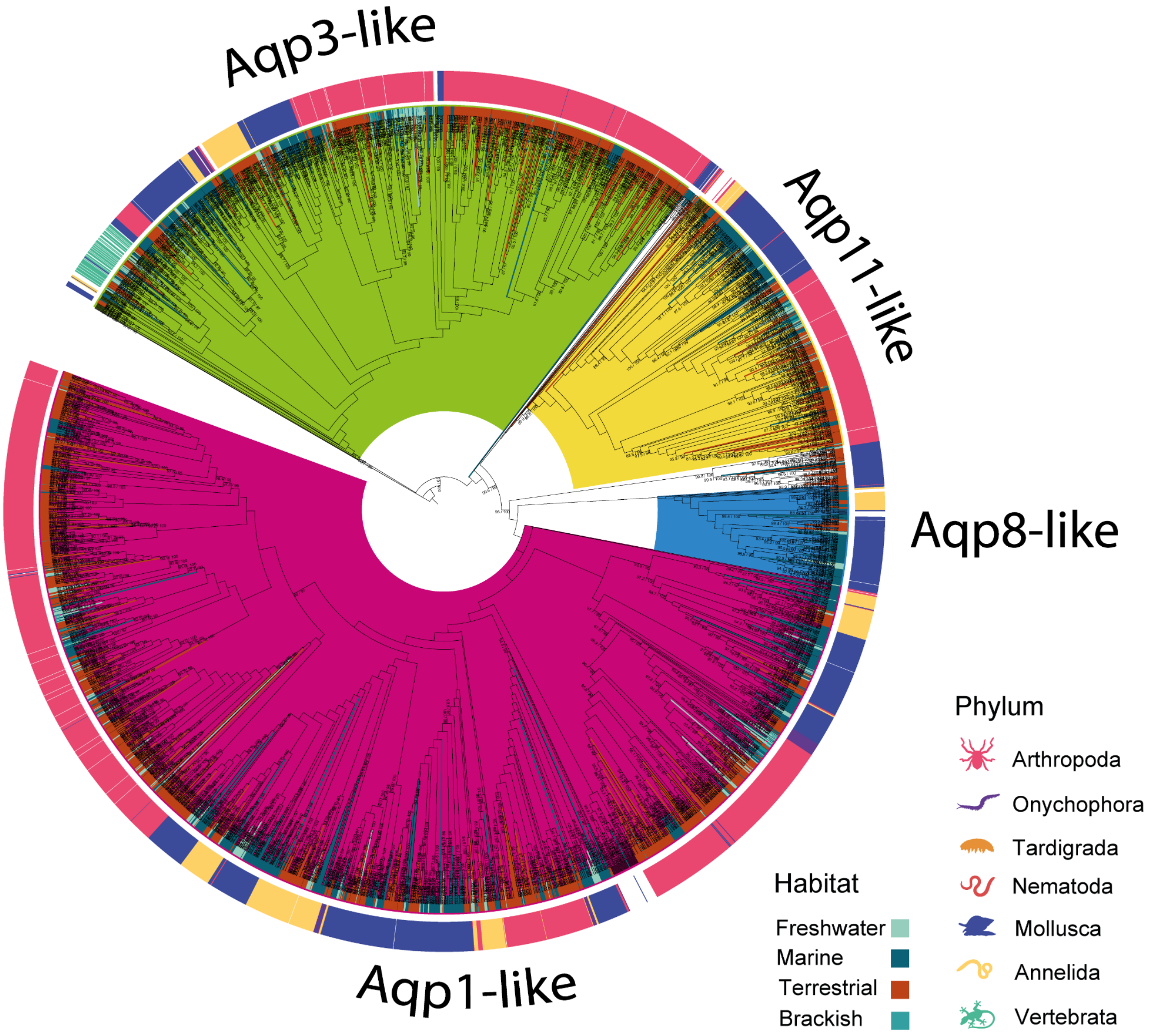

**Figure S2. Phylogenetic tree of 481 Aqp11-like plus 27 backbone sequences in the 7 studied animal phyla.**

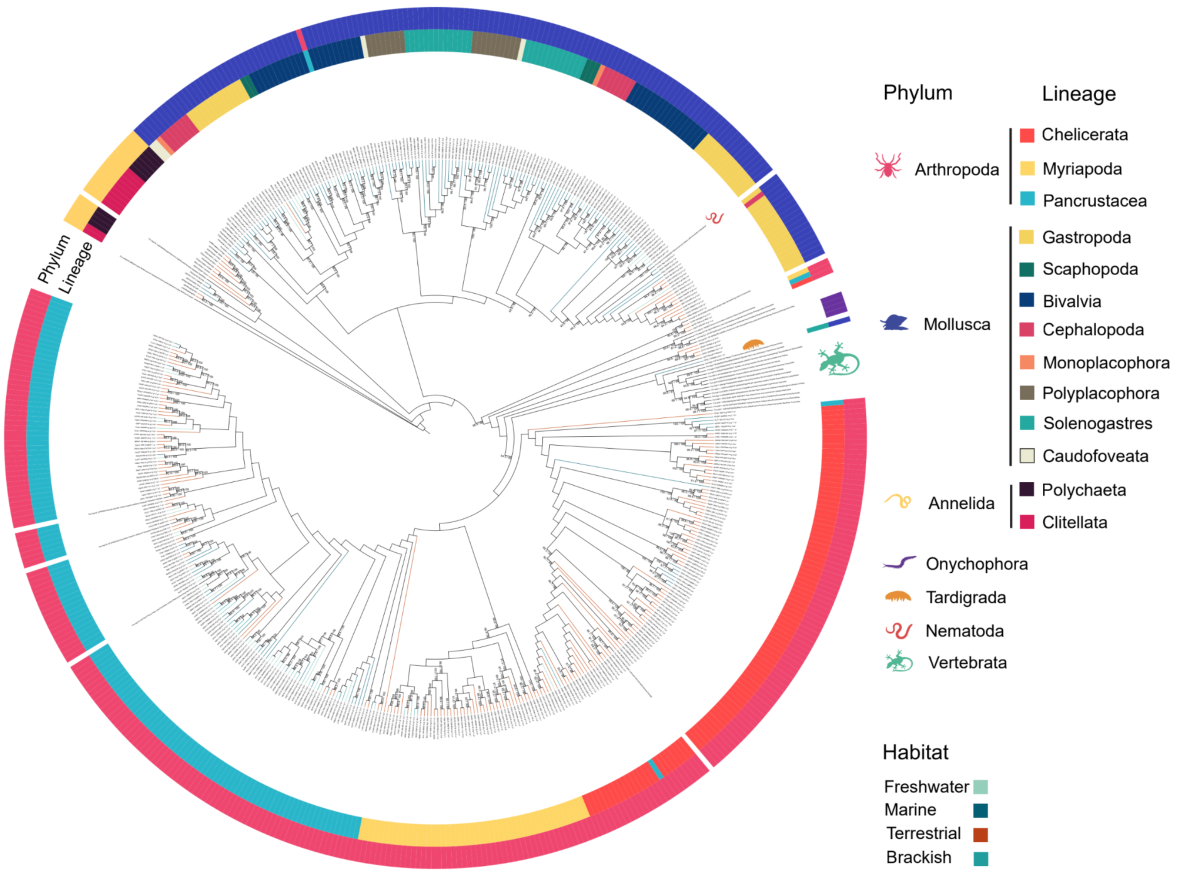

**Figure S3. Uncollapsed consensus phylogenetic tree of 2,202 Aqp1-like plus 63 backbone sequences in 6 studied phyla.**

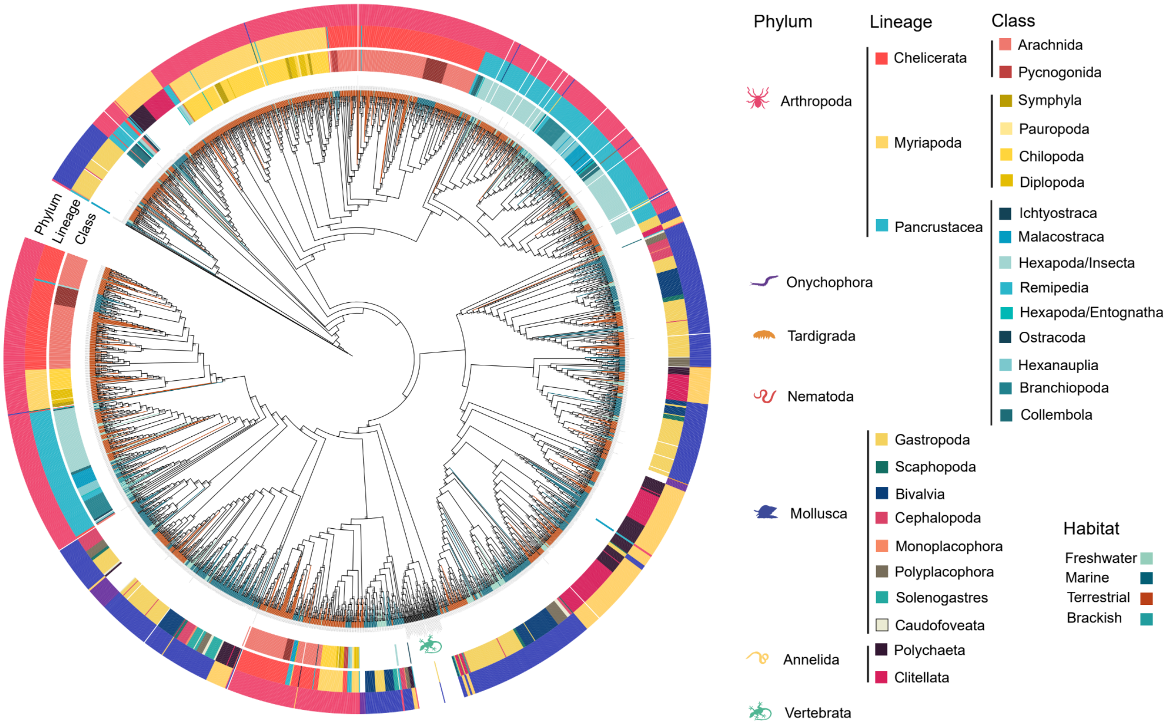

**Figure S4. Consensus phylogenetic tree of 1,156 arthropod Aqp1-like plus 64 backbone sequences.**

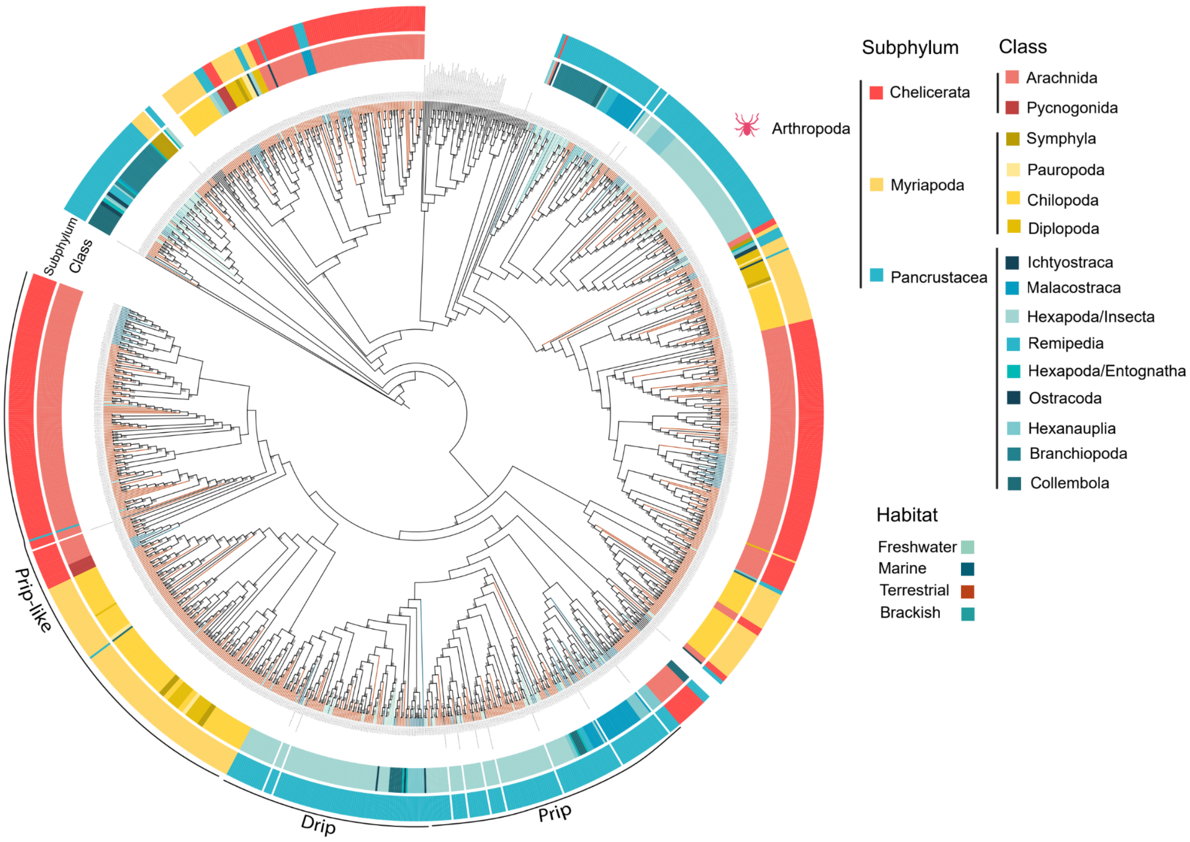

**Figure S5. Phylogenetic tree of 162 Aqp8-like plus 19 backbone sequences in 4 studied animal phyla.**

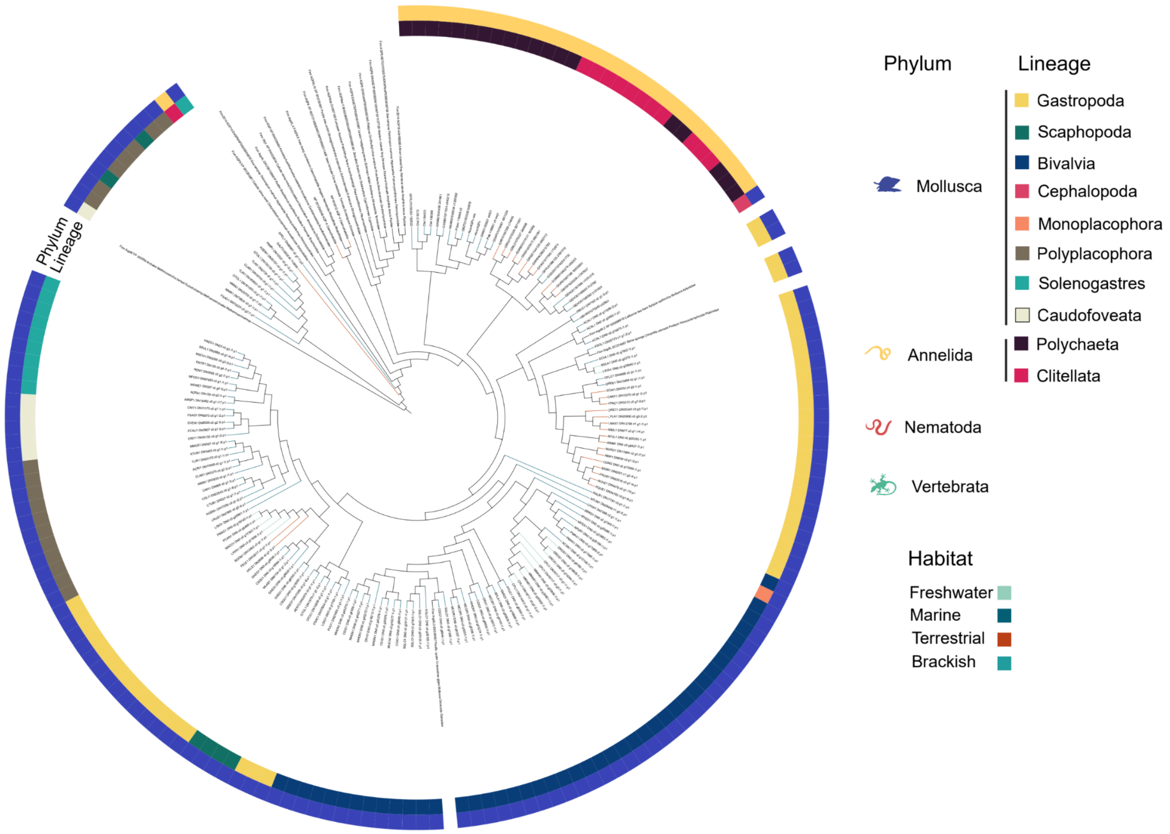

**Figure S6. Uncollapsed consensus phylogenetic tree of 1,108 Aqp3-like plus 68 backbone sequences in the 7 studied animal phyla.**

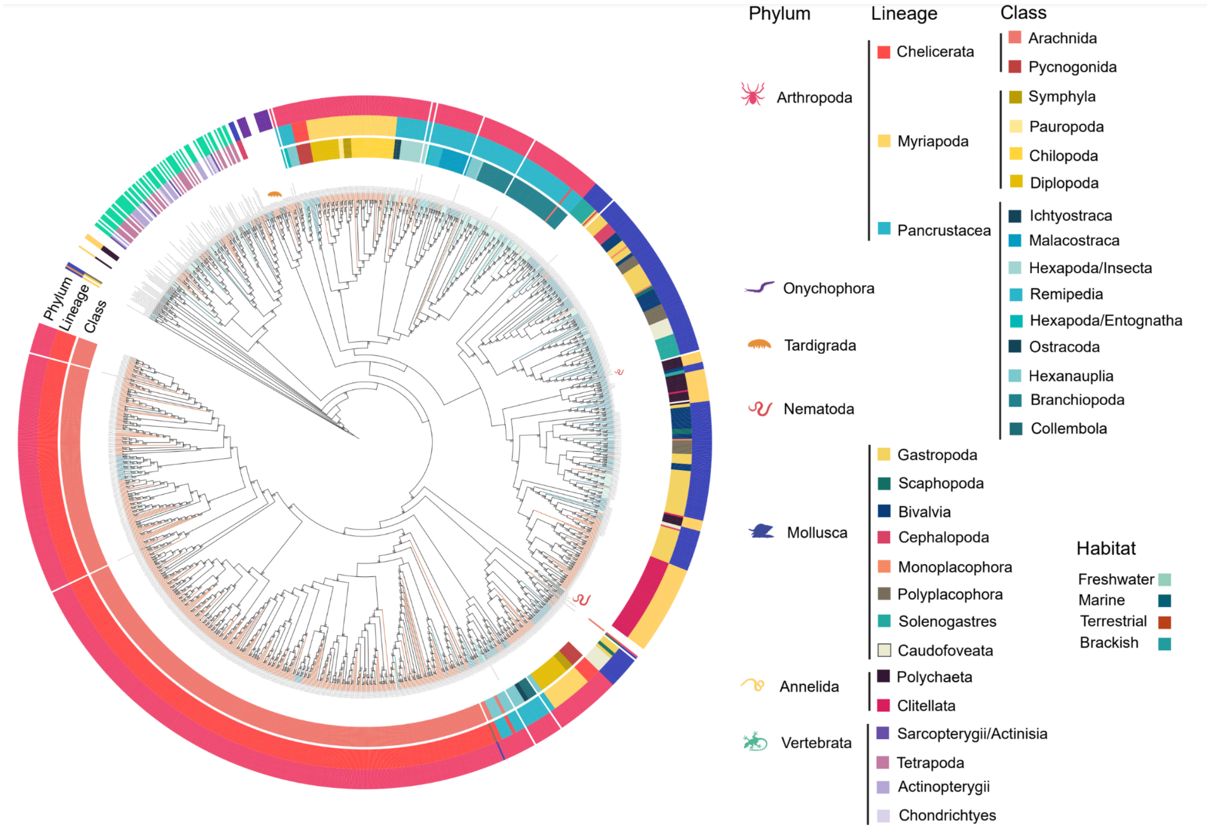

**Figure S7. Phylogenetic tree of 176 Eglps plus 7 backbone sequences in the insects included.**

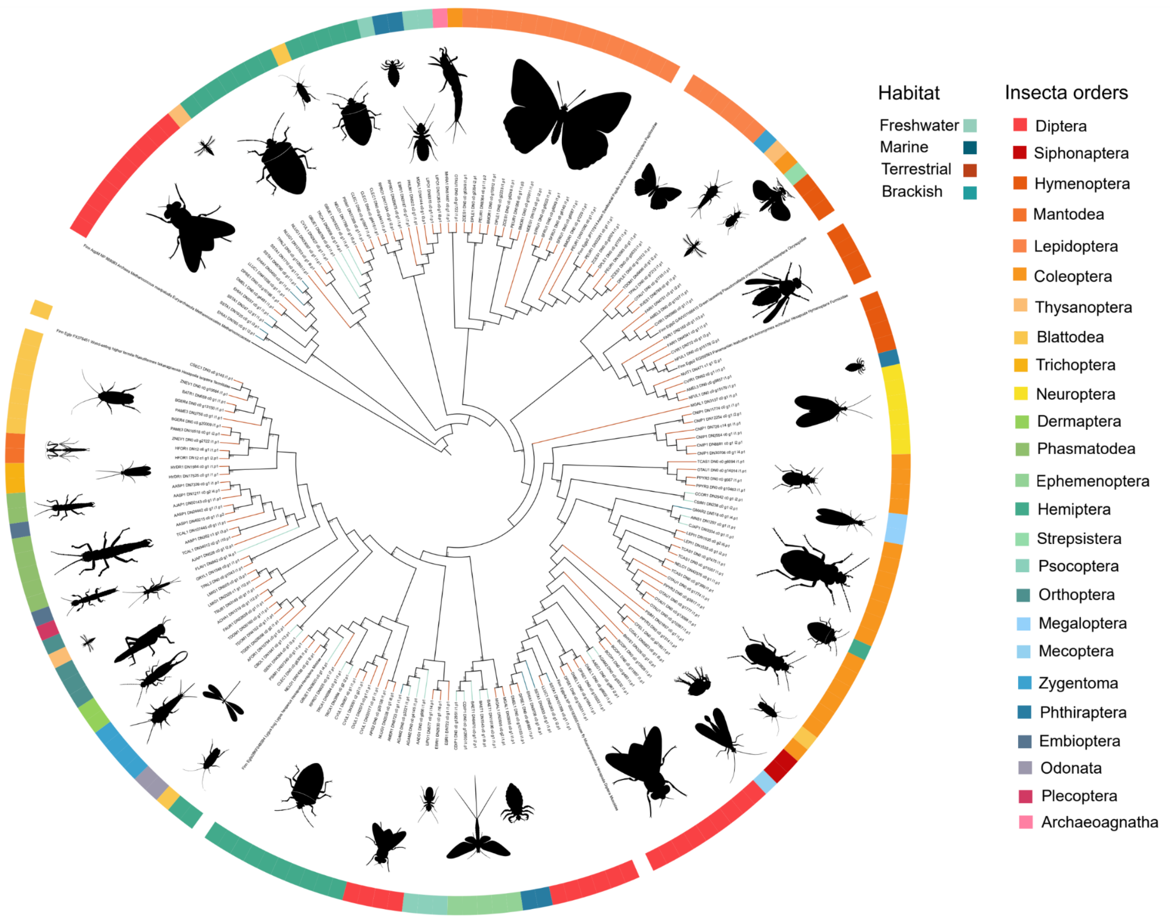

**Figure S8. ML phylogenies built for all Aqp1-like and all arthropod Aqps to test the effect of alignment trimming using trimAL.** Outer circumference represents the phylum and inner circumference represents the lineage inside each phylum. Different trimming options affect the formation of Aqp clades, with more extensive trimming options generating clades formed by Aqps of mixed origins. **A,B.** Without applying any trimming step. **C,E.** trimAL automated1 option, which opted for the strict option in both cases based on sequence number and identity score. **D,F.** trimAL gappyout option, which automatically selects a gap threshold (-gt) based on gap distribution. **G.** Removing all sequences that have a gap in arthropod Aqps. Applying this option to Aqp1-like alignment produced the same alignment as the one without any type of trimming. **H-M.** Selecting 3 different gap thresholds (1 - fraction of sequences with a gap allowed).

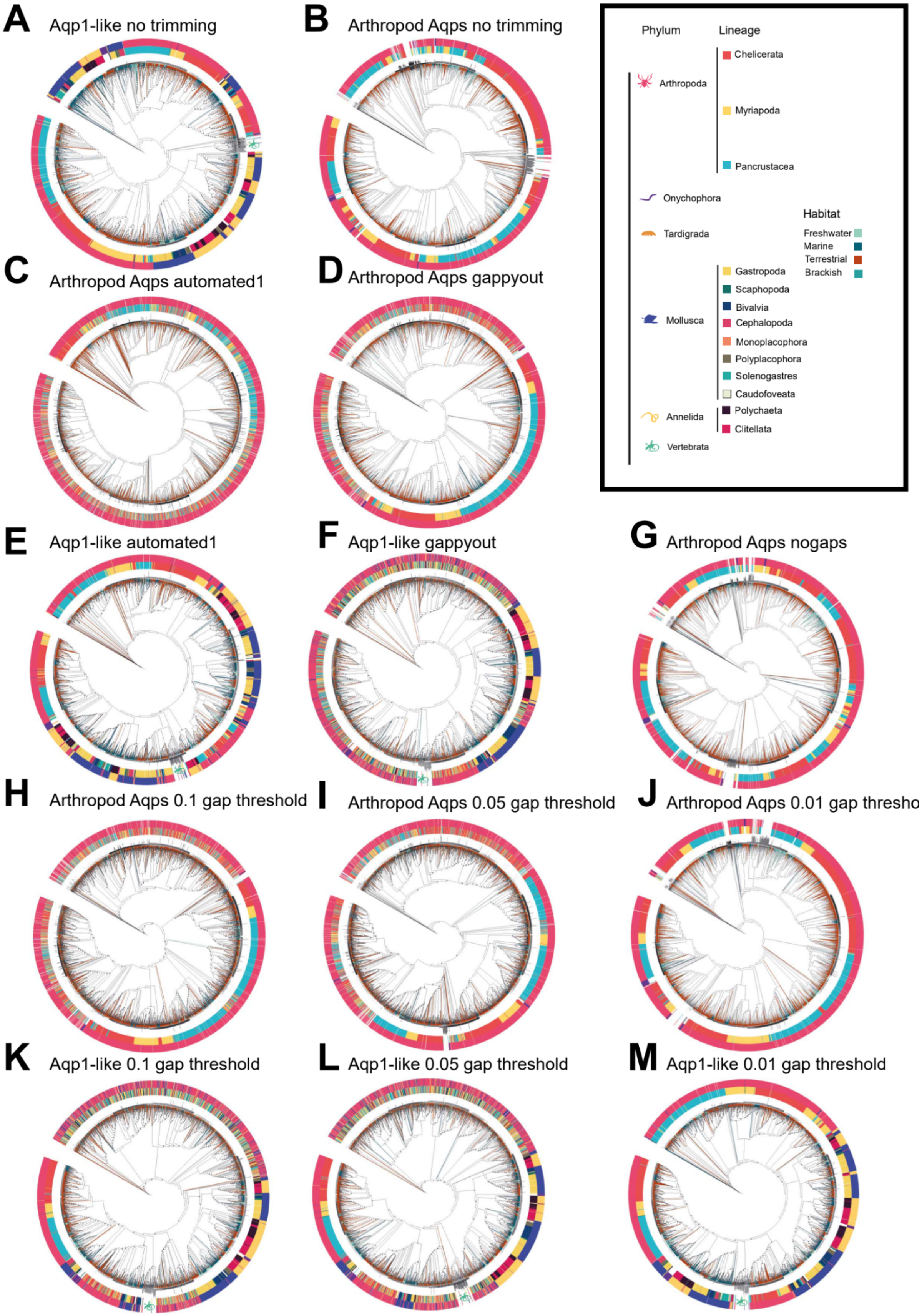
A Aqp1-like no trimming
B Arthropod Aqps no trimming
Phylum
Lineage
Arthropoda
Chelicerata
Myriapoda
Pancrustacea
Onychophora
Tardigrada
Habitat
Freshwater
Marine
Terrestrial
Brackish
Mollusca
Gastropoda
Scaphopoda
Bivalvia
Cephalopoda
Monoplacophora
Polyplacophora
Solenogastres
Caudofoveata
Annelida
Polychaeta
Clitellata
Vertebrata
C Arthropod Aqps automated1
D Arthropod Aqps gappyout
E Aqp1-like automated1
F Aqp1-like gappyout
G Arthropod Aqps nogaps
H Arthropod Aqps 0.1 gap threshold
I Arthropod Aqps 0.05 gap threshold
J Arthropod Aqps 0.01 gap thresho
K Aqp1-like 0.1 gap threshold
L Aqp1-like 0.05 gap threshold
M Aqp1-like 0.01 gap threshold
